# Microglia-to-neuron signaling increases lipid droplet metabolism, enhancing neuronal network activity

**DOI:** 10.1101/2025.08.03.668224

**Authors:** Ana P. Verduzco Espinoza, Na Na, Loraine Campanati, Priscilla Ngo, Kristin K. Baldwin, Hollis T. Cline

## Abstract

Microglia regulate neuronal circuit plasticity. Disrupting their homeostatic function has detrimental effects on neuronal circuit health. Neuroinflammation contributes to the onset and progression of neurodegenerative diseases, including Alzheimer’s disease (AD), with several microglial activation genes linked to increased risk for these conditions. Inflammatory microglia alter neuronal excitability, inducing metabolic strain. Interestingly, expression of *APOE4*, the strongest genetic risk factor for AD, affects both microglial activation and neuronal excitability, highlighting the interplay between lipid metabolism, inflammation, and neuronal function. It remains unclear how microglial inflammatory state is conveyed to neurons to affect circuit function and whether *APOE4* expression alters this intercellular communication. Here, we use a reductionist model of human iPSC-derived microglial and neuronal monocultures to dissect how the *APOE* genotype in each cell-type independently contributes to microglial regulation of neuronal activity during inflammation. Conditioned media (CM) from LPS-stimulated microglia increased neuronal network activity, assessed by calcium imaging, with *APOE4* microglial CM driving higher neuronal firing rates than *APOE3* CM. Both *APOE3* and *APOE4* neurons increase network activity in response to CM treatments, while *APOE4* neurons uniquely increase presynaptic puncta with *APOE4* microglial CM. CM-derived exosomes from LPS-stimulated microglia can mediate increases to network activity. Lastly, increased network activity is accompanied by increased lipid droplet (LD) metabolism and blocking LD metabolism abolishes network activity. These findings illuminate how microglia-to-neuron communication drives inflammation-induced changes in neuronal circuit function, demonstrate a role for neuronal LDs in network activity, and support a potential mechanism through which *APOE4* increases neuronal excitability.

## Introduction

Microglia perform essential roles in neuronal circuit function and synaptic plasticity, mediating synaptic pruning and modulating synaptic strength (1–3). As the brain’s resident immune cells, they are also the main drivers of neuroinflammation, a phenomenon implicated in neurodegenerative diseases, including Alzheimer’s disease (AD). Microglia are activated in response to pathogens, injury, and other insults, altering their gene expression and function to recruit relevant cells to resolve the inflammatory process. Although the extent to which this disruption in microglial homeostatic function affects neuronal health is not fully understood, multiple genome-wide association studies have linked microglial activation genes, including *TREM2*, *APOE*, *BIN1*, and *SPI1*, with an increased risk of AD (4–7). Thus, understanding the mechanisms by which microglial inflammatory responses alter neuronal function is essential for better understanding the early mechanisms underlying neurodegeneration.

Inflammatory microglia alter neuronal network activity and metabolism. In vivo treatment with lipopolysaccharide (LPS), a commonly used model of inflammation, can increase neuronal excitability, induce seizures, and—upon prolonged exposure—lead to oxidative stress, synapse dysfunction, and cell death (8, 9). Additionally, as microglia adopt an inflammatory state, they undergo a metabolic shift that impacts surrounding cells. Neuronal metabolism specifically is altered by inflammatory microglia, as neurons are known to have an altered rate of respiration and they can switch to alternate energy sources to sustain network activity (10, 11). Identifying how microglial inflammatory responses are communicated to neurons and how they affect neuronal metabolism is critical for understanding the vulnerabilities contributing to neuroinflammation-driven neurodegeneration.

A key mechanism that could mediate microglia-to-neuron communication is the release of exosomes. Exosomes are membrane-bound particles that contain a variety of functional cargoes, including proteins, RNAs, and metabolites, which change based on donor cell type and state (12–14). Microglia, like most cells, secrete exosomes under basal conditions, but inflammatory microglia secrete exosomes with distinct cargoes that can propagate the inflammatory phenotype to recipient microglia (15–18). Exosomes from inflammatory microglia have also been shown to inhibit neuroprogenitor proliferation (19) and potentially impair cognitive function, as blocking exosome release in an LPS mouse model improved behavioral outcomes (20). Although their role in regulating circuit function is not fully understood, microglial exosomes are promising candidates for relaying inflammatory cues to alter neuronal excitability.

Among the genes associated with increased risk for AD, *APOE* stands out for its strong influence on both microglial reactivity and neuronal function. *APOE*, which encodes the Apolipoprotein E lipid carrier, has three allelic variants of which *APOE4* is the strongest genetic risk factor for late-onset AD (21). In microglia, *APOE4* expression drives a pro-inflammatory transcriptional profile, characterized by lower expression of homeostatic genes and higher expression of inflammatory cytokine genes (22–26). Additionally, *APOE4* microglia accumulate lipid droplets (LDs)—a hallmark of their inflammatory response— even in the absence of a pathological stimulus and they can induce tau phosphorylation, apoptosis, and disrupt neuronal network activity (22, 27). Interestingly, recent work demonstrated that *APOE4*-related neuronal pathologies improved in the absence of microglia in an AD mouse model, suggesting that microglial *APOE4* expression could drive neurodegeneration (28). In addition to the effects of *APOE4* expression in microglia, neuronal *APOE4* expression is also associated with increases in neuronal excitability (25, 29, 30), posing the question of whether neuronal *APOE* genotype could influence their susceptibility to excitotoxicity. However, it remains unclear how APOE4 expression in each cell type independently contributes to microglia-to-neuron signaling during inflammation, a question that is difficult to address *in vivo*, where multiple cell types contribute to the inflammatory response.

In this study, we investigate microglia-to-neuron signaling in the context of inflammation. Using induced pluripotent stem cell (iPSC)-derived microglial and neuronal monocultures, we find that exosomes from inflammatory microglia increase synchronized neuronal network activity. Assessing the bioactivity of microglial conditioned media (CM) shows that CM from LPS-stimulated inflammatory microglia similarly increases neuronal network activity. We use CRISPR editing to dissect the contributions of APOE3 and isoAPOE4 genotypes in microglial to neuron signaling. Signaling from inflammatory isoAPOE4 microglia drives increased network activity in neurons, regardless of neuronal *APOE* genotype, while *APOE4* neurons uniquely increase presynaptic puncta in response to *APOE4* microglial CM. Considering the cellular energy source for the increased neuronal activity, we find that CM from inflammatory microglia decrease neuronal lipid droplet load. We identify neuronal LD metabolism as a mechanism that is required to sustain network activity and that adapts with microglia-driven hyperexcitability.

## Results

### Generating and characterizing iPSC-derived iNs and inflammatory iMGLs

To enable our study of how *APOE* genotype influences microglial-to-neuron signaling during inflammation, we CRISPR-edited an established *APOE3/E3* iPSC line (31) to change amino acid residue 112 from cysteine to arginine and generate isogenic *APOE4/E4* iPSCs (isoAPOE4; Fig. S1A). iPSC lines were karyotyped and characterized via a single nucleotide polymorphism (SNP) array to confirm euploidy and they were evaluated for their expression of pluripotency markers TRA 1-60 and SSEA4 by flow cytometry (Fig. S1B-C). Expression of pluripotency markers Nanog, SSEA-4, SOX2 and Oct-4 was also confirmed in both iPSC lines via immunocytochemistry (Fig. S1D-E).

Induced neurons (iNs) were generated following a Classic NGN2 induction protocol (Fig. S2A) (32) with modifications to enhance neuronal maturation (Enhanced NGN2 neural induction; Fig. S2B) (33). As previously reported, by day 24, iNs generated with a Classic NGN2 induction express *APOE*, pan neuronal markers *MAP2* and TUBB3, and excitatory synapse markers *VGLUT1* and *VGLUT2* at higher levels than inhibitory markers like *GAD1* and *GAD2*, which we confirmed by RT-qPCR (Fig. S2A, F). To enhance neuronal maturation, we used a prolonged 49-day protocol with additional neurotrophic factors (Fig. S2B). Immunocytochemistry revealed that like iNs generated with a Classic NGN2 induction, iNs generated via an Enhanced NGN2 induction protocol express excitatory synaptic markers vGlut1 and 2 while lacking expression of inhibitory marker Gad67 (Fig. S2E). We further characterized iNs via immunocytochemistry to evaluate neuronal marker NeuN and cortical marker Cux1 protein expression (Fig. S2C). With a Classic NGN2 induction of 24 days (Fig. S2A), 86.7% of APOE3 and 89% of isoAPOE4 iNs express NeuN, and 72.9% and 67.3% of NeuN-expressing cells also express cortical marker Cux1 (Fig. S2D). Compared to Classic NGN2 D24 iNs, however, Enhanced D49 iNs have greater proportions of both NeuN-expressing cells. NeuN+ cells increase significantly to 94.4% and 94.1% of APOE3 and isoAPOE4 iNs, respectively, with Cux1+ cells averaging 93.8% and 88.8% of NeuN+ cells (Fig. S2D). Indicative of their improved maturity, Enhanced D49 iNs also display spontaneous network activity, which we assessed via live imaging with calcium indicator Fluo-4AM (Movie S1), a feature not seen in Classic D24 iNs.

To generate induced microglia-like cells (iMGLs), we followed an established protocol in which iPSCs are first differentiated into CD43-expressing hematopoietic progenitors (HPCs) and further differentiated into mature microglia over the course of 7 weeks (34) (Fig. S3A). The expression of CD43 in HPCs was characterized via immunocytochemistry, and we found 93.7% and 92.5% of HPCs express CD43 in APOE3 and isoAPOE4 cell lines, respectively, with little variation across batches (SEM = 1.4 and 1.6; Fig. S3B). Mature iMGLs were functionally validated via a phagocytosis assay with fluorescent beads. In APOE3 iMGL cultures, 89% of cells displayed phagocytic activity, while isoAPOE4 iMGL cultures had a significantly larger proportion of phagocytic cells at 95% (Fig. S3C). We observed via immunocytochemistry that a large proportion of both APOE3 and isoAPOE4 iMGLs express microglial markers PU.1 (93.4% and 97.9%, respectively) and Iba1 (89.3% and 89.7%, respectively) with no significant difference between *APOE* genotypes (Fig. S3D). Immunocytochemistry also confirmed that APOE3 and isoAPOE4 iMGLs express microglial markers Trem2 and P2RY12 (Fig. S3E). Via RT-qPCR, we observed similar expression levels of *TREM2*, *IBA1*, *P2RY12*, and *APOE* in APOE3 compared to isoAPOE4 iMGLs (Fig. S3G).

Given previous reports of *APOE4* microglia having a disrupted homeostatic state and enhanced inflammatory response compared to *APOE3* microglia (22, 27, 35), we wanted to evaluate iMGL signaling under basal and inflammatory conditions. To induce a robust inflammatory response in iMGLs, we treated cells with 50ng/mL LPS every other day for a week, allowing exosomes and other signaling factors produced as part of the inflammatory response to accumulate in the conditioned media (CM; Fig. 1A). Compared to PBS, LPS-treatment caused clear morphological changes, including hypertrophic cell bodies, and increased expression of inflammatory cytokine genes *IL6*, *IL10* and *CCL2* evaluated by RT-qPCR in both APOE3 and isoAPOE4 iMGLs (Fig. S3G). In APOE3 iMGLs, LPS treatment decreased expression of homeostatic *P2RY12*; however, in isoAPOE4 iMGLs, LPS did not affect *P2RY12* expression, suggesting an altered inflammatory response (Fig. S3G). LPS-treatment also caused accumulation of LDs in both APOE3 and isoAPOE4 iMGLs compared to PBS controls, but isoAPOE4 iMGLs had a significantly greater LD load than their APOE3 counterparts both with PBS- and LPS-treatments (Figure S3F). We assessed the expression of genes involved in LD metabolism and found no differences in *PLIN2*, *ACAT1*, or *DDHD2* expression between APOE3 and isoAPOE4 iMGL with or without LPS treatment (Fig. 3SG). We did, however, observe differences in the expression of *DGAT-1*, which encodes an enzyme involved in the synthesis of triacylglycerol (TAG), a major component of LDs. Surprisingly, *DGAT-1* expression was significantly lower in isoAPOE4 iMGLs compared to APOE3 iMGLs and, whereas *DGAT-1* expression decreased with LPS treatment in APOE3 iMGLs, it did not change in isoAPOE4 iMGLs treated with LPS (Fig. 3SG). This could reflect a compensatory mechanism in iMGLs with high LD load (36). Overall, these data show that a week-long LPS treatment successfully induces a robust inflammatory response in both APOE3 and isoAPOE4 iMGLs, with isoAPOE4 iMGLs having a distinct inflammatory phenotype that could alter downstream intercellular signaling.

**Figure 1.**
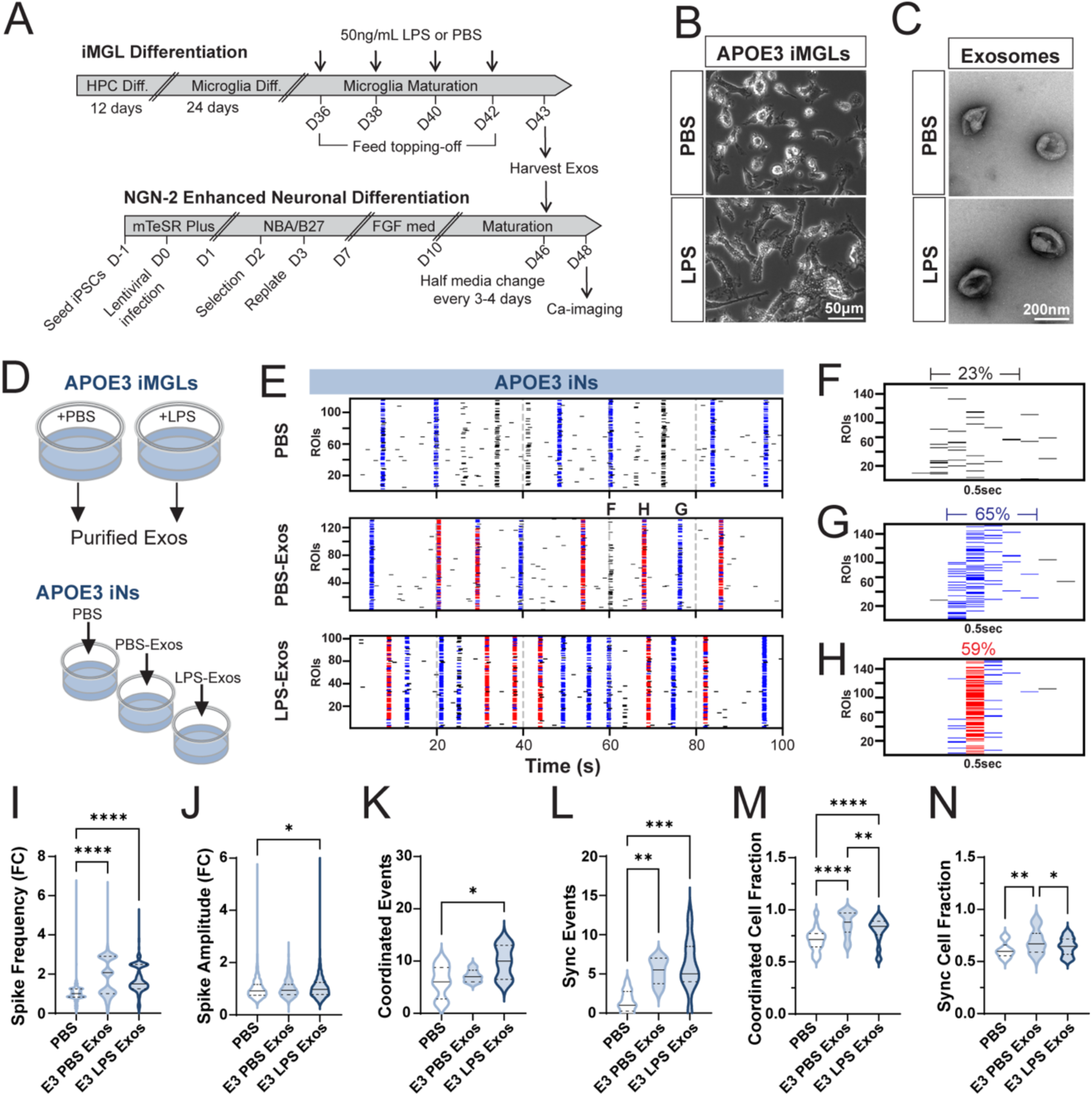
Exosomes from inflammatory iMGLs increase iN network activity. A) Differentiation timelines for iMGLs and iNs. B) Representative brightfield images of D43 iMGLs after a 7-day treatment with PBS or 50ng/mL LPS. C) Transmission electron micrographs of iMGL exosomes. D) Schematic of treatment conditions. Exosomes were isolated from APOE3 iMGL conditioned media (CM) via sequential ultracentrifugation, and APOE3 D46 iNs received either PBS or exosomes from PBS- or LPS-treated iMGLs (PBS-Exos and LPS-Exos, respectively). E) Representative raster plots of Ca-imaging recordings in APOE3 iNs at D48. Imaging with calcium indicator Fluo4-AM was conducted after 48hr treatment with PBS or iMGL Exos. Blue tick marks represent coordinated events, defined as instances where >50% of cells spiked within a 250ms window (5 frames). Red tick marks represent synchronized events, defined as instances where >50% of cells spiked within a 50ms window (1 frame). F-H) Enlarged examples of activity bursts over 0.5 sec. F) Correlated activity where only 23% of cells spiked within a 250ms window, not meeting criteria for a coordinated event. G) Coordinated event in which 65% of cells spiked within a 250ms window. H) Synchronous event in which 59% of cells fired within a 50ms window. I-J) Violin plots of spike frequency (I) and amplitude (J) per cell normalized as the fold change relative to the mean of PBS-treated iNs. (N = 665-1132 cells from 2 independent experiments). K-L) Violin plots showing number of coordinated events (K) and synchronized events (L) per 100 sec recording. (N = 6-12 recordings, 1-3 recordings per well from 2 independent experiments). M-N) Violin plots showing the fraction of cells engaged in each coordinated event (M; N = 43-89 coordinated events from 2 independent experiments) or synchronized event (N; N = 18-53 synchronized events from 2 independent experiments). Violin plot lines represent median (solid) and quartiles (dashed). Median and quartile values can be found in Table S4. One-way ANOVA with Tukey test for multiple comparisons. * p < 0.05, ** p < 0.01, *** p < 0.001, and **** p < 0.0001.

### Exosomes from inflammatory microglia increase neuronal network activity

To assess the contribution of exosomes in microglia-to-neuron signaling in the context of inflammation, we harvested exosomes from PBS- and LPS-treated APOE3 iMGLs (Fig. 1A-B). iMGL exosomes were isolated from CM via sequential ultracentrifugation (Fig. S4A), as we previously described (37), and their characteristic cup-like morphology was identified by transmission electron microscopy (Fig. 1C). Additionally, we demonstrated by Western blot that iMGL exosomes contain exosomal markers Alix and Flotillin-1 (Fig. S4B).

We treated D46 APOE3 iNs with either PBS (vehicle) or purified exosomes from PBS- or LPS-treated iMGLs (Fig. 1D, PBS, PBS-exos, LPS-exos). Exosome samples were normalized based on a protein assay, and equal amounts were administered across conditions. After 48hrs, we imaged spontaneous neuronal activity in iN cultures over 100 second epochs using the calcium indicator Fluo4-AM (Fig. 1E). Assessing individual neuron firing rates, we found that APOE3 neurons significantly increased spike frequency when treated with either PBS- or LPS-exosomes compared to the control treatment with PBS alone (Fig. 1I). Additionally, spike amplitude increased with LPS-exosomes compared to PBS controls (Fig. 1J).

iN cultures exhibited repeated bouts of network activity throughout the 100 sec epochs, with varying proportions of neurons engaged in each bout (Fig. 1E, Movies S1-3). This led us to wonder whether exosomes, which increased the neuronal firing rate, could also increase network connectivity, increasing the frequency of—and the proportion of cells recruited to—network events. To address this question, we quantified two types of network activity: coordinated and synchronized events (Fig. 1F-H). We defined coordinated events as instances where >50% of neurons in the field of view (FOV) spiked within a broad 250ms window, representing the temporally dispersed network activity (Fig. 1G). We then defined synchronized events as the subset of coordinated events in which >50% of neurons in the FOV were co-active within a shorter 50msec window, representing only the temporally precise activity of neuronal networks with robust synaptic connectivity (Fig. 1H).

We quantified the incidence of coordinated (Fig. 1K) and synchronized events (Fig. 1L) as well as the fraction of cells active in both types of network events (Fig. 1M-N). Compared to treatment with PBS, treatment with PBS-exosomes did not affect the incidence of coordinated events in iN cultures (Fig. 1K) but rather increased the synchronicity of coordinated activity resulting in a greater number of synchronized events (Fig. 1L). PBS-exosome treatment in iN cultures also promoted the recruitment of additional neurons into both coordinated and synchronized events, increasing the fractions of co-active cells compared to control PBS treatment (Fig. 1M-N). Compared to PBS treatment, treatment with LPS-exosomes increased the incidence of global coordinated events and enhanced synchronicity in coordinated networks, increasing also the numbers of synchronized events in iN cultures (Fig. K-L). LPS-exosomes, however, recruited significantly fewer neurons to coordinated events than PBS-exosomes and did not increase the fraction of co-active cells in synchronized events compared to control PBS treatment (Fig. 1M-N). This suggests that while PBS-exosomes enhance the synchronicity of existing coordinated activity and expand synchronized neuronal networks, LPS-exosomes primarily increase the incidence of coordinated and synchronized events without promoting the recruitment of additional neurons into these networks.

In summary, APOE3 iMGL exosomes increase neuronal spiking frequency independent of microglial inflammatory status. PBS- and LPS-exosomes, however, have distinct effects on network activity. While PBS-exosomes appear to foster circuit refinement, LPS-exosomes drive a more generalized increase in the activity of both strongly and weakly connected neuronal networks.

### Inflammatory *APOE4* microglial signaling drives greater network activity than *APOE3*

We next sought to evaluate how microglial and neuronal *APOE* genotype affects microglia-to-neuron signaling via conditioned media (CM) in the context of inflammation. Using both APOE3 and isoAPOE4 iN cultures, we performed a half-media exchange on iNs with conditioned media from PBS- and LPS-treated APOE3 and isoAPOE4 iMGLs (Fig. 2A). To control for the dilution of neuronal media, we used PBS- or LPS-unconditioned media (UCM) as control treatments. After 48hrs, we imaged spontaneous neuronal activity in APOE3 iN and isoAPOE4 iN cultures (Fig. 2B, G). Treating APOE3 and isoAPOE4 iNs with either PBC UCM or LPS UCM showed comparable low baseline activity in iNs of both genotypes, which is likely due to the dilution of neurotrophic factors in neuronal media. Treatment with CM from PBS-treated APOE3 iMGLs decreased spike amplitude in APOE3 iNs while increasing spike amplitude in isoAPOE4 iNs (Fig. 2C, H). CM from PBS-treated isoAPOE4 iMGLs, however, did not affect spike amplitude in iNs of either *APOE* genotype. CM from LPS-treated APOE3 and isoAPOE4 iMGLs increased spike amplitude in both APOE3 and isoAPOE4 iNs, with isoAPOE4 LPS CM causing a significantly greater spike amplitude than APOE3 LPS CM (Fig. 2C, H).

**Figure 2.**
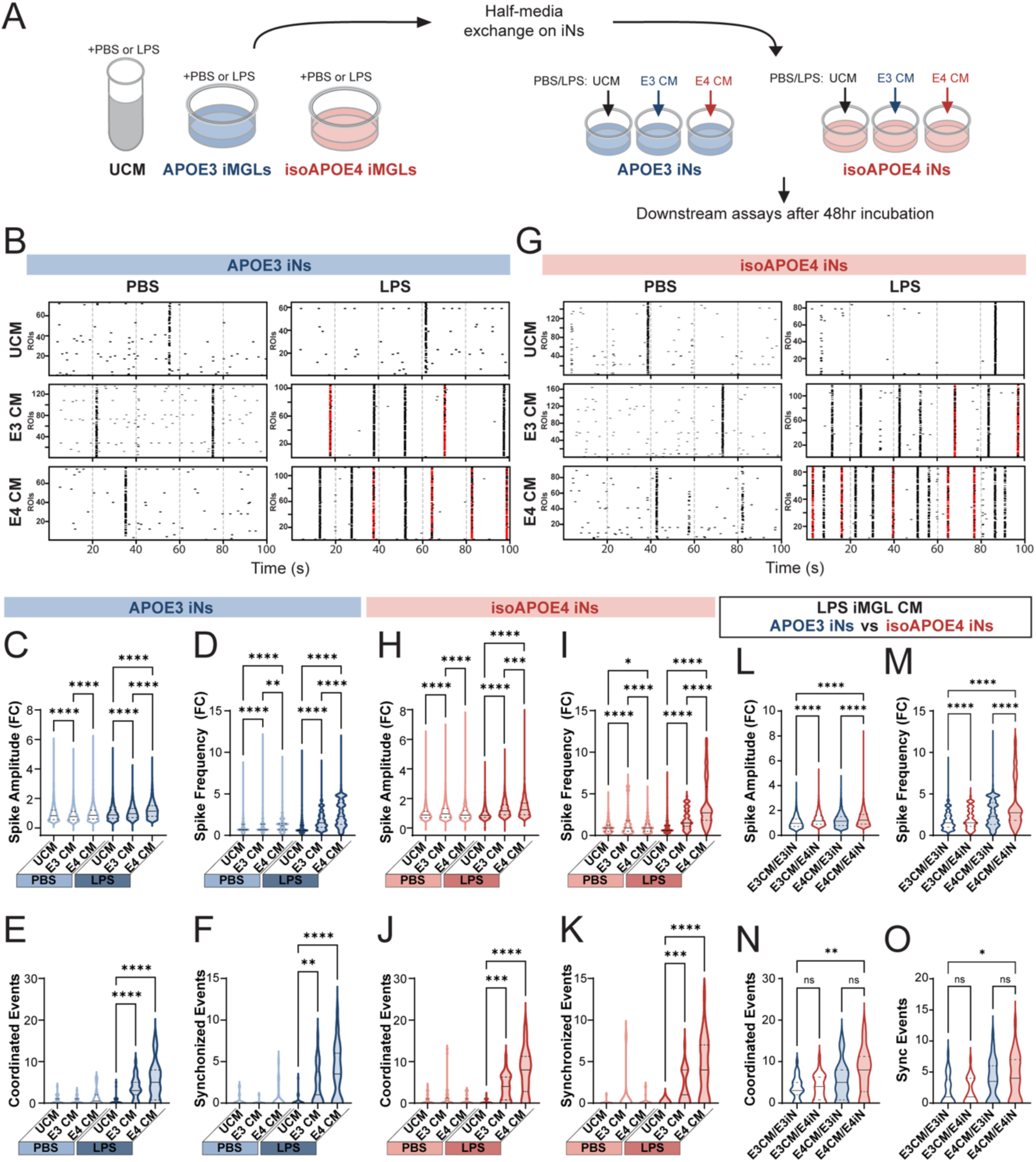
Inflammatory *APOE4* microglial signaling drives greater neuronal network activity than *APOE3.* A) Schematic of experimental design. APOE3 (B-F, blue) and isogenic APOE4 (G-K, red) iNs received a half-media exchange with either unconditioned media (UCM) or iMGL CM and underwent Ca-imaging with Fluo4-AM 48hrs later. B,G) Representative raster plots of Ca-imaging recordings from APOE3 (B) and isogenic APOE4 (G) neurons treated with UCM or CM from PBS- or LPS-treated iMGLs. Red tick marks represent synchronized events. C-D,H-I) Violin plots of spike amplitude (C,H) and spike frequency (D,I) plotted as fold change relative to the average of PBS UCM-treated iNs (N = 1223-3211 cells from 4 independent experiments; Statistical comparisons were performed separately within the PBS-treated and LPS-treated groups using one-way ANOVAs followed by Tukey tests). E-F,J-K) Violin plots showing number of coordinated events (E,J) and synchronized events (F,K) per 100 sec recording (N = 26-33 recordings, 1-2 recordings per well from 4 independent experiments; Statistical comparisons were performed separately within the PBS-treated and LPS-treated groups using Kruskal-Wallis tests with Dunn’s multiple comparison tests). L-O) Violin plots showing statistical comparisons in spike frequency (L), spike amplitude (M), coordinated event count (N), and synchronized event count (O) across neuronal *APOE* genotype for iNs treated with LPS-stimulated iMGL CM. For L-O, statistical comparisons were performed with one-way ANOVA with Sidak multiple comparisons test. Violin plot lines represent median (solid) and quartiles (dashed), and values can be found in Table S4. * p < 0.05, ** p < 0.01, *** p < 0.001, and **** p < 0.0001.

LPS-treated iMGL CM showed a genotype-dependent effect with isoAPOE4 CM driving a greater increase in neuronal activity than APOE3 CM (Fig. 2D, I). In addition, LPS-CM increased both coordinated and synchronized events in iNs independent of their APOE genotype (Fig. 2E-F, J-K) and this increase in coordinated events occurs by recruitment of neurons into the highly co-active networks (Fig. S5B, D). By contrast, CM from PBS-treated iMGLs increased spike frequency in iNs independent of the genotypes of the iMGLs or the iNs but had no impact on coordinated or synchronized events (Fig. 2D-F, I-K). Overall, these data indicate that inflammatory iMGL CM has a distinct effect on neuronal network activity and synchronicity.

We performed additional post-hoc comparisons to evaluate whether neuronal *APOE4* genotype further increases neuronal activity in response to CM from inflammatory iMGLs. When treated with either APOE3 or isoAPOE4 LPS iMGL CM, isoAPOE4 iNs showed significantly higher spike amplitude and spike frequency compared to APOE3 iNs (Fig. 2L-M). IsoAPOE4 iNs treated with isoAPOE4 LPS iMGL CM significantly increased both coordinated and synchronized events compared to APOE3 iNs treated with APOE3 LPS iMGL CM. We did not observe a difference in coordinated or synchronized event count between APOE3 and isoAPOE4 iNs when receiving the same LPS iMGL CM treatment (Fig. 2N-O). These findings suggest that *APOE4* genotype in neurons increases neuronal excitability in response to CM but *APOE4* genotype in microglia is responsible for driving increased network activity during inflammation.

Increased network activity could result from structural changes to synapses. We immunolabeled iNs for presynaptic marker synapsin and quantified the number of puncta (Fig. S5E-F). This showed that isoAPOE4 iNs increased synapsin puncta following treatment with both PBS and LPS isoAPOE4 iMGL CM (Fig. S5F, right) but APOE3 iNs were not affected (Fig. S5F, left) These data suggest that isoAPOE4 iNs exhibit greater synaptic plasticity than their APOE3 counterparts and that isoAPOE4 iMGL CM might have a unique ability to remodel presynaptic elements in isoAPOE4 iNs. Additionally, these findings show that structural changes to presynaptic elements are not required to increase network activity, suggesting that other mechanisms, such as increased intrinsic excitability may contribute to greater network activity. Overall, our findings suggest that microglial *APOE* genotype modulates their signaling capabilities, especially under inflammatory conditions. Interestingly, although we detected similar changes to network activity in both APOE3 and isoAPOE4 iNs in response to these treatment conditions, the underlying structural mechanisms mediating these changes in activity may depend on neuronal *APOE* genotype.

### Increased iN network activity is accompanied by a depletion of neuronal LDs

Given the robust increase in neuronal network activity that we observed, and its known energetic cost, we questioned how neurons could fuel such sudden changes in energy demand. In many cell types, LDs can serve as energy stores and the TAGs that make up LDs can be metabolized to fuel a variety of cellular processes (38). Although neurons are not thought to rely on lipids as an energy source, a recent study demonstrated that neurons can utilize LD-derived free fatty acids (FFAs) to sustain synaptic activity (39). We investigated whether the conditions that increased neuronal network activity affected LD load in neurons by staining CM-treated D48 iNs with BODIPY-493/503 (Fig. 3A). To capture LDs in the entirety of neuronal cell bodies, we quantified LDs using maximum intensity projections of 10μm confocal z-stacks acquired in 1μm steps and normalized LDs to nuclei count. We found that in APOE3 iNs, isoAPOE4 PBS iMGL CM decreased LDs compared to PBS UCM (Fig. 3B, left) while in APOE4 iNs neither APOE3 nor isoAPOE4 PBS CM altered neuronal LD load (Fig. 3B, right). Compared to LPS UCM, treatment with both APOE3 and isoAPOE4 LPS iMGL CM significantly decreased LDs in iN cultures of both *APOE* genotypes (Fig. 3B). To rule out potential artifacts arising from the 10μm projections, we performed alternate analysis and quantified LDs in single confocal planes and normalized LD count to the area of βIII-tubulin-positive neurites. We found similar, although smaller, decreases in LD load with this alternate analysis (Fig. S6A-B). Interestingly, the LPS-CM treatments which decreased neuronal LD load are the same conditions that increased network activity, supporting our hypothesis that LD metabolism might help neurons meet the energy demand driven by increased network activity.

**Figure 3.**
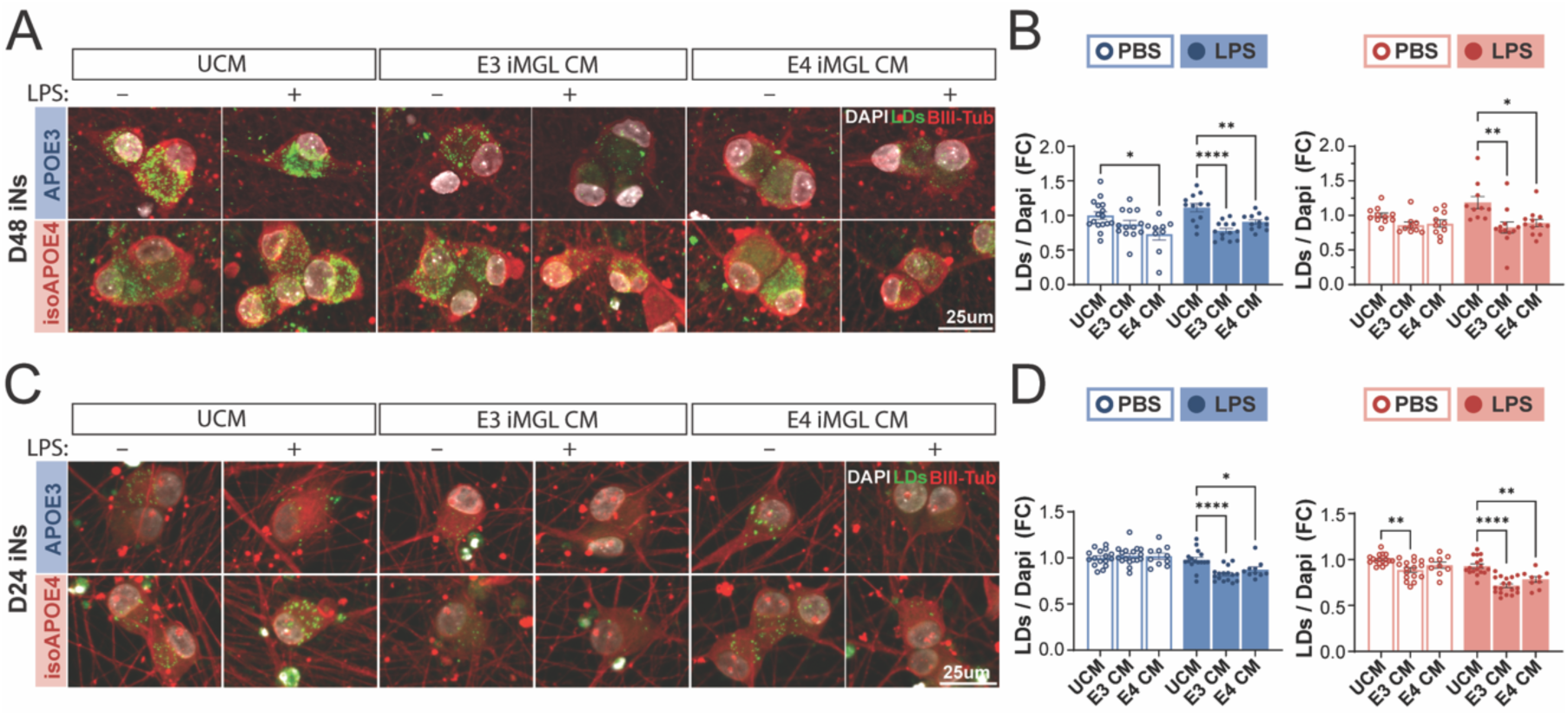
Inflammatory iMGL conditioned media decreases neuronal lipid droplet load. D48 or D24. iNs were treated with unconditioned media (UCM) or conditioned media (CM) from PBS- or LPS-treated iMGLs for 48hrs and immunolabeled for βIII-tubulin (red). Lipid droplets (LDs) were labeled with BODIPY-493/503 (green) and nuclei were labeled with DAPI (white). A) Representative images of UCM or CM-treated APOE3 (top, blue) and isoAPOE4 (bottom, red) D48 iNs. Images are max intensity projections of 10μm confocal z-stacks taken at 1μm steps and cropped for visualization. B) Quantification of LDs in APOE3 (left, blue) and isoAPOE4 (right, red) iNs. LDs were normalized to DAPI-labeled nuclei per field of view, averaged by well, and graphed as the fold change relative to UCM PBS conditions (N=9-16 wells from 3-4 independent experiments). C) Representative images of CM-treated D24 iNs and D) quantification of LDs/nuclei (N = 9-17 wells from 2-3 independent experiments). Statistical comparisons were performed separately within the PBS-treated and LPS-treated groups using one-way ANOVAs followed by Tukey’s post hoc tests. Bar graphs represent the mean with SEM. * *p* < 0.05, ** *p* < 0.01, *** *p* < 0.001, and **** *p* < 0.0001.

We then tested whether changes in LD load were present in less mature neuronal cultures in which circuit activity is less robust. For this, we used the Classic NGN2 induction method (32) with minor modifications (Fig. S2A). We treated D24 iNs with the same iMGL CM conditions described above and used BOIDPY-493/503 to label and quantify LDs (Fig. 3C-D). While D24 iNs had fewer LDs than the more mature D48 iNs (Fig. S6C), they had a similar response to the iMGL CM treatments. In APOE3 iNs, CM-treatment from APOE3 and isoAPOE4 PBS-iMGLs did not alter LD load, while in isoAPOE4 iNs, CM-treatment from APOE3 PBS-iMGLs decreased LDs (Fig. 3D). Like more mature D48 iNs, D24 iNs of both APOE3 and isoAPOE4 genotypes had significantly decreased LDs when treated with CM from LPS-treated iMGLs, regardless of iMGL *APOE* genotype (Fig. 3D). This suggests that inflammatory microglia can modulate neuronal LD metabolism in iN cultures at different maturation stages, and this modulation is not dependent on pre-existing iN network activity. Overall, these results support our hypothesis that neuronal LD stores may be metabolized to support the increased energy demand of changes to spontaneous activity.

### LD metabolism is required to sustain network activity

To further investigate LD metabolism in iN cultures, we employed an additional pair of APOE3 and isoAPOE4 iPSC lines to ensure that our observations were not specific to a single isogenic background (Fig. S7). The additional iPSC lines appeared genetically normal after characterization with SNP array, although karyotyping showed that 25% of APOE3 iPSCs (5/20 cells tested) had trisomy of chromosome 1 (Fig. S7A-B). Both APOE3 and isoAPOE4 iPSC lines showed a high level of expression of pluripotency markers TRA 1-60 and SSEA-4 by FACS and expressed Nanog, SSEA-4, SOX2, and Oct-4 by immunocytochemistry (Fig. S7A-D). We also confirmed that D24 iNs generated from both iPSC lines express neuronal markers NeuN, Cux1, vGlut1 and vGlut2 by immunocytochemistry (Fig. S7E-G), and confirmed the expression of *APOE*, *MAP2*, and *TUBB3* as well as synaptic markers like *VGLUT1* and *VGLUT2* via RT-qPCR (Fig. S7H).

To investigate whether lipid droplets are actively metabolized in iNs under basal conditions, we treated iN cultures with KLH45, an inhibitor of neuron-specific lipase DDHD2 which breaks down LD TGs into FFAs that can be used as energy (40). We hypothesized that if iNs continuously metabolize LDs under basal conditions to support functions such as synaptic activity, blocking LD metabolism would result in a rapid accumulation of LDs (Fig. 4A). We conducted a time-point experiment where we treated APOE3 and isoAPOE4 D24 iNs with 5μM KLH45 or vehicle (EtOH) for 2, 8, or 24hrs. Strikingly, we found that iNs treated with KLH45 rapidly accumulate LDs. Within 8hrs of exposure to KLH45, iNs show more than a 2-fold increase in number of LDs, and this accumulation further increases by 24hrs (Fig. S8A-D). Interestingly, both APOE3 and isoAPOE4 D24 iNs accumulated LDs at similar rates. When we evaluated the expression of genes involved in LD metabolism, there was also no difference in the expression levels of *DDHD2*, *PLIN-1*, *DGAT-1*, and *ATGL* between APOE3 and isoAPOE4 iNs (Fig. S2F, S6H).

**Figure 4.**
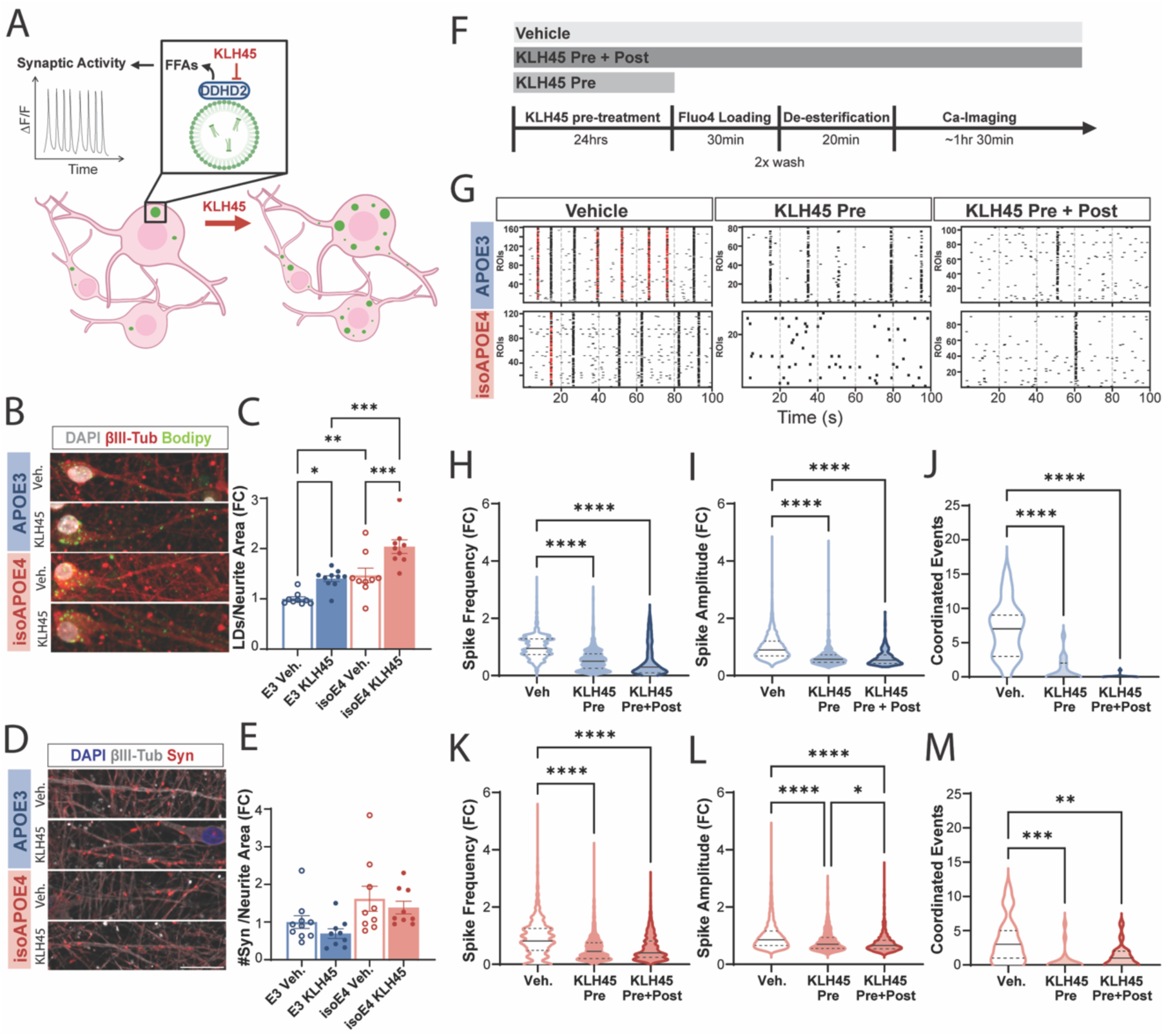
LD metabolism is required for neuronal network activity. A) Schematic of KLH45 mechanism of action and our experimental hypothesis. DDHD2 sits on the surface of lipid droplets (LDs) and breaks down triglycerides (TGs) into free fatty acids (FFAs). KLH45 inhibits DDHD2, which in turn results in cells accumulating LDs. We hypothesized that if LD metabolism supports neuronal network activity, KLH45 would result in a decrease of network activity. B-E) D48 iNs were treated with vehicle or 5μM KLH45 for 24hrs and analyzed for relative changes in LDs and synapses. B) Representative images of D48 iNs immunolabeled for βIII-tubulin (red) with LDs labeled with BODIPY-493/503 (green) and nuclei labeled with DAPI (white). Images are max intensity projections of 10μm confocal z-stacks taken at 1μm steps and cropped for visualization. C) Quantification of LDs per field of view, normalized to neurite area, and averaged by well. Data are shown as fold change relative to the APOE3 Vehicle condition (N = 9-10 wells from 2 independent experiments; Two-way ANOVA with Sidak correction for multiple comparisons). D) Representative images of iNs immunolabeled for βIII-tubulin (white) and synapsin 1/2 (red) with nuclei labeled with DAPI (blue). Confocal images were cropped for visualization. E) Quantification of synapsin puncta per field of view, normalized to neurite area, and averaged by well. Data are shown as fold change relative to the APOE3 Vehicle condition (N = 9-10 wells from 2 independent experiments; Two-way ANOVA with Sidak correction for multiple comparisons). F) Schematic of KLH45 treatment timeline used for Ca-imaging with Fluo4-AM in D48 iNs. iNs received a 24hr pre-treatment with either vehicle (EtOH) or 5μM KLH45. KLH45 was either removed (KLH45 Pre) or kept (KLH45 Pre+Post) during Ca-imaging. G) Representative raster plots. Red tick marks represent synchronized events. H-M) Quantification of Ca-imaging features for APOE3 (H-J, blue) and isoAPOE4 (K-M, red) iNs. H,K) Quantification of average spike frequency per cell normalized as the fold change to the average of Veh-treated iNs (N = 326-3236 cells from 2 independent experiments). I,L) Quantification of average spike amplitude per cell, normalized as the fold change to the average of Veh-treated iNs (N = 326-3236 cells from 2 independent experiments). J,M) Violin plots showing quantification of coordinated events per 100sec recording session (N = 17-20 recordings, 1-2 recordings per well from 2 independent experiments). One-way ANOVA with Tukey correction for multiple comparisons. Bar graphs represent the mean and error bars represent SEM. Violin plot lines represent median (solid) and quartiles (dashed), and values can be found in Table S4. * *p* < 0.05, ** *p* < 0.01, *** *p* < 0.001, and **** *p* < 0.0001.

We then looked at D48 Enhanced iNs and found that a 24hr KLH45 treatment also results in an accumulation of LDs (Fig. 4B-C, S8G-H). Interestingly, isoAPOE4 iNs had a higher abundance of LDs compared to APOE3 iNs both with vehicle and KLH45 treatment, a difference we did not observe in D24 iNs. This suggests that neuronal *APOE* genotype could affect neuronal LD metabolism in more mature iN cultures.

To test whether LD metabolism is required to sustain spontaneous neuronal network activity, we performed calcium imaging on D48 iNs pre-treated with vehicle or KLH45 for 24hrs, with the drug treatment either maintained or removed during the calcium imaging (Fig. 4F-G). In both APOE3 and isoAPOE4 iNs, we saw a marked decrease in spike frequency and amplitude with KLH45 pre-treatment, regardless of whether the drug was maintained during the imaging period (Fig. 4H-I, K-L). KLH45 also caused a stark decline in coordinated events (Fig. 4J, M). Interestingly, changes in neuronal activity were not accompanied by changes in the number of pre-synaptic synapsin puncta (Fig. 4D-E). These results were validated in our second set of APOE3 and isoAPOE4 cell lines (Fig. S8I-Q). To rule out the possibility of the decline in neuronal activity being caused by KLH45-induced toxicity, we used confocal imaging to quantify healthy, non-condensed nuclei based on size and DAPI signal intensity. We found no changes in the number of healthy nuclei per field of view (Fig. S8E-F). Overall, these findings support our hypothesis that LD metabolism is required to sustain spontaneous network activity in iN cultures, and the decrease in activity driven by blocking LD metabolism is not preceded by detectable structural changes to pre-synaptic compartments.

## Discussion

In this study, we show that microglia drive neuronal activity through secreted factors, including exosomes, and that they specifically enhance network activity under inflammatory conditions. We demonstrate that microglial *APOE4* expression further increases neuronal excitability and show that neuronal LD metabolism is essential to sustain spontaneous network activity. These findings provide a mechanistic link between inflammation, neuronal excitability, and lipid metabolism, all of which are implicated in Alzheimer’s disease.

While CM from resting microglia drives an increase in spike frequency, CM from LPS-stimulated microglia not only causes a greater increase in spike frequency but also enhances synchronous network activity in iN cultures (Fig. 2). Notably, exosomes isolated from microglial CM recapitulate these signaling bioactivities; while both PBS- and LPS-exosomes increase spike frequency, only LPS-exosomes enhance global network activity (Fig. 1E-K), positioning exosomes as potential mediators of this microglia-neuron regulation of activity. While previous work has shown that LPS-induced inflammation can increase neuronal activity, most of these studies were done *in vivo* and in acute brain slices, which contain astrocytes and other cell types that contribute to the inflammatory response (9, 41, 42). Our reductionist approach allowed us to show that signaling from microglia alone can directly alter neuronal activity during LPS-induced inflammation. Interestingly, prior work on microglia derived microvesicles (MVs)—which differ from exosomes in cellular origin and cargo—showed that both resting and LPS-stimulated MVs increased neuronal excitability to similar extents (43). A direct comparison of microglial exosomes and MVs, as well as an assessment of exosome- and MV-depleted CM, is needed to clarify the specific contributions of exosomes in microglia-to-neuron signaling.

Our reductionist experimental design also allowed us to investigate the independent contributions of microglial and neuronal APOE genotype to microglial modulation of neuronal activity. Given the known influence of *APOE4* genotype on microglial reactivity (22–24, 26) and our observed role of microglia exosome signaling in modulating network activity, we investigated how microglial *APOE4* genotype affects this modulation. Resting *APOE4* microglia had more LDs than their *APOE3* counterparts, and this difference was exacerbated upon LPS-stimulation (Fig. S3F). This aligns with other reports of *APOE4* genotype increasing microglial reactivity and LD accumulation (22, 25, 27, 36, 44). When evaluating CM bioactivity, we found that resting *APOE4* microglial CM increases neuronal activity after the 2-day treatment compared to *APOE3*. We further observed that *APOE4* amplifies the inflammatory response, with LPS-treated *APOE4* microglia driving a greater increase in network activity than *APOE3* (Fig. 2). Thus, our findings suggest that microglial *APOE4* can drive neuronal hyperexcitability via intercellular signaling, particularly during inflammation. Notably, a recent study in iPSC-derived microglia found a 7-day treatment with *APOE4* resting microglial CM reduced network activity in neurons (27). Given that we observed a striking depletion of LDs accompanying the increased network activity with a 2-day CM treatment, and that blocking LD metabolism abolished neuronal activity, we speculate that prolonged CM exposure may deplete neuronal LDs, impairing their ability to sustain network activity. More work is needed to understand the timescale at which microglial CM regulates neuronal activity and the underlying mechanisms that mediates this modulation. Additionally, other microglial activation pathways, including purinergic signaling, are known to also alter neuronal network activity and synchronicity (45). Whether the *APOE4* genotype affects microglia-neuron signaling in the context of additional activation pathways merits further investigation.

Neuronal *APOE4* expression is associated with neuronal hyperexcitability preceding neurodegeneration in diseases like AD (46). A variety of mechanisms have been implicated in this effect, including neuronal *APOE4* driving increased synaptic protein expression and excitatory activity (25) and decreased inhibitory tone in brain regions vulnerable to degeneration (29, 47). Recent work also suggests that neuron-specific *APOE* genotype drives this excitation-inhibition imbalance (29). Although we show that microglial *APOE4* genotype drives an increase in neuronal activity, we did not observe consistent evidence of neuronal *APOE* genotype affecting excitability. While PBS-treated *APOE4* neurons had increased network activity compared to *APOE3* neurons (Fig. 1), we did not see the same trend in the control conditions of media exchange (Fig. 2) and KLH45 treatment experiments (Fig. 4). We also observed similar increases in activity in both *APOE3* and *APOE4* neurons upon treatment with microglial CM (Fig. 2). Although we detected increased synapsin puncta following treatment with APOE4 microglial CM, this was not correlated with increased network activity. Specifically, unlike *APOE3* neurons which had no change to presynaptic elements in response to microglial CM, *APOE4* neurons increased presynaptic puncta when treated with *APOE4* microglial CM. This suggests that neuronal APOE could be affecting the underlying mechanisms that give rise to hyperexcitability in other models. Given that several lines of work show that certain neuron sub-populations are more susceptible to *APOE* and AD driven hyperexcitability (46, 47), more work is needed to characterize the extent to which *APOE* genotype affects neuronal function in models of susceptible neuronal subtypes.

It is well established that neuronal activity is energetically costly and that increases in synaptic activity drive adaptive changes to neuronal metabolism (48, 49). While the direct contribution of LD metabolism in fueling neuronal activity is unclear, there is a growing body of evidence supporting a critical role for LD metabolism in maintaining neuronal health. Loss-of-function mutations in the neuron-specific TAG lipase *DDHD2*, for example, cause spastic paraplegia, a neurodegenerative disorder characterized by massive neuronal LD accumulation, limb weakness, and intellectual disability (40, 50, 51). In our NGN2-induced neurons, we found LDs in both 24- and 48-day-old cultures, with older neurons displaying a greater LD load (Fig. 3). Upon treatment with LPS-stimulated microglial CM, which increased neuronal activity, we observed a reduction in LD abundance—suggesting upregulated LD metabolism (Fig. 3). Given the recent reports demonstrating that neuronal activity drives DDHD2-mediated FFA synthesis (52) and that neuronal LD-derived FFAs can be used as oxidative phosphorylation fuel during periods of heightened activity (39), we hypothesized that our iNs might use LDs to fuel both spontaneous and CM-driven activity. We probed LD dynamics by inhibiting the LD lipase DDHD2 using KLH45. In young (D24) iNs, KLH45 treatment led to a significant accumulation of LDs within just 8hrs, indicating ongoing LD turnover even in the absence of network-wide activity (Fig. S8A-B). This LD turnover could support low-level activity or membrane remodeling in these maturing neurons, as LD triglyceride metabolism has been linked to axon regeneration in models of nerve crush (53). In more mature (D48) iNs, which exhibit robust network activity, KLH45 treatment for 24hrs led to an accumulation of LDs and concurrent suppression of network activity (Fig. 4G-M). Thus, our data show that DDHD2-mediated LD metabolism is required to sustain spontaneous network activity. We propose that neuronal LDs serve as energy reservoirs supporting neuronal activity, and that blocking their metabolism impairs this function. Since neurons are particularly sensitive to oxidative stress from fatty acid catabolism (54), future studies investigating how neuronal LD metabolism is regulated and whether reliance on this pathway contributes to neurodegeneration under conditions of inflammation, *APOE4* expression, and heightened energy demand would be informative.

## Acknowledgments

We thank all Cline lab members for their advice and feedback. We acknowledge the expert assistance of Scott Henderson, Kimberly Vanderpool, and Theresa Fassel of The Core Microscopy Facility at The Scripps Research Institute. We thank Katherine Spencer and Shakib Omari at the Dorris Neuroscience Center Core Microscopy facility for their support and guidance. This work was supported by grants from the National Institutes of Health (1RF1AG079517-01) and a Fellowship from the Helen Dorris Foundation to A.V.E.

## Materials and Methods

### Generation of isogenic APOE4/E4 human induced pluripotent cell (iPSC) lines

We used two previously generated and characterized independent Wellderly APOE3/E3 iPSC lines, HE463#7 and HE19#1 (1). These lines are available on WiCell in a collection called Topol Lab’s Next Gen Cell Lines with the following accession numbers: HE463#7 (SCRP2307i) and HE0019#1 (SCRP2101i). APOE3/E3 iPSC lines were genetically modified using the CRISPR/cas9 system to change the amino acid at position 112 from a cysteine to an arginine, generating two isogenic APOE4/E4 iPSC lines. We performed CRISPR/cas9 editing following the protocol described in Lin et al., 2018 with minor modifications. In brief, we used a sgRNA sequence (5’-CCTCGCCGCGGTACTGCACC-3’) within ten nucleotides from the target site to amino acid 112 and cloned it into pSpCas9-2A-GFP plasmid (Addgene #48138). We also used single-strand oligodeoxynucleotides (ssODN) to convert APOE3→APOE4 while introducing a silent mutation at the protospacer adjacent motif (PAM) site to prevent recurrent Cas9 cutting (5’GAGGAGACGCGGGCACGGCTGTCCAAGGAGCTGCAGGCGGCGCAGGCCCGGCTGGGCGCGGACATGGAGGACGTGCGCGGCCGGCTGGTGCAGTACCGCGGCGAGGTGCAGGCCATGCTCGGCCAGAGCACCGAGGAGCTGCGGGTGCGCCTCGCCTCCCACCTGCGCAAGCTGC GTAAG-3’). We dissociated iPSCs in accutase (Stem Cell Technologies, 7920) with 10μM Y-27632 (Stem Cell Technologies, 72304) and electroporated cells with 7.5μg of CRISPR/cas9 sgRNA and 15μg of ssODN using an Amaxa nucleofector unit and Human Stem Cell Nucleofector Kit 1, according to manufacturer’s instructions (Lonza, VPH-5012). Cells were then seeded onto Matrigel-coated plates (Corning, 354277; Greiner Bio-One, 657160) in mTeSR Plus (Stem Cell Technologies, 100-0276) with 10μM Y-27632. 48hrs after the transfection, we performed FACS to isolate GFP-expressing cells and seeded them on Matrigel coated plates.

When cells reached ∼70% confluency (5-7 days), we split them at low density and collected a portion of the cell pool to isolate genomic DNA and amplify the target region using PCR. PCR products were cleaned up using a kit (Qiagen, 28104) and sent for Sanger sequencing. We looked for evidence of editing using Benchling alignment software. When evidence of editing was found, cells were dissociated with accutase and 5x10^3^ cells were plated in 10cm dishes in mTeSR with 10μM Y-27632. After 5-7 days, single colonies were manually picked under a dissecting microscope, each seeded into a well on a 12-well plate (Greiner Bio-One, 665102). When clones reached ∼70% confluency, they were passaged, and half of cells were collected for sequencing. Successfully edited clones were then split into 6-well plates to do a second round of sequencing to confirm APOE4/E4 genotype and further characterization.

### Culture and characterization of iPSCs

Two pairs of APOE3/E3 (HE19#1 and HE463#7) and isogenic APOE4/E4 (HE19#1 E34#44 and HE463#7 E34#17) iPSC lines were used in this study. iPSCs were thawed in mTeSR Plus with 10μM Y-Y-27632 and seeded on 6-well dishes coated with Matrigel. For maintenance, iPSCs were fed every 1-3 days, depending on confluency, and passaged 1-2 times per week as small clusters using 1μM EDTA (Invitrogen, 15575-020). 1X Penicillin-Streptomycin was added to all culture media (Life Technologies, 15140-122).

We used immunofluorescent labeling and flow cytometry to characterize the expression of pluripotency markers SSEA4 and TRA-1-60 in all lines. In brief, iPSCs were dissociated with accutase and fixed in suspension with 4% paraformaldehyde (EMS, 50980487) for 20min. Approximately 250,000 fixed cells per sample were incubated in blocking buffer (5% heat inactivated FBS in PBS) for 30 minutes at room temperature and then incubated for 20 min in 5% Alexa Fluor 488 anti-human TRA-1-60-R and APC anti-human SSEA-4 antibodies (1:20, BioLegend, 330613 and 330417) in wash buffer (1% heat inactivated FBS in PBS). Cells were washed twice, resuspended in wash buffer, and analyzed by flow cytometry (Bio-Rad ZE5 Cell Analyzer). Gates were drawn based on unlabeled controls using the FLowJo software (BD Biosciences Version 10.10.0).

We also verified the expression of pluripotency markers Nanog, Oct4, Sox2, and SSEA-4 via immunocytochemistry (see “Immunofluorescent labeling”) and confocal microscopy. For this, iPSCs were grown on Matrigel-coated glass-bottom 96-well plates (Cellvis, NC0536760). To rule out chromosomal abnormalities, all iPSC lines were characterized via single nucleotide polymorphism (SNP) array (UCSD IGM Genomics Center). Genomic DNA was extracted using a GeneJET Genomic DNA Purification kit following manufacturer’s instructions (Thermo Fisher Scientific, K0721) and 1μ of DNA was submitted for analysis. iPSCs were also tested for mycoplasma using a kit (ATCC, 30-1012K).

### Lentivirus generation

We generated lentiviral particles to transduce iPSCs for neuron differentiation using human embryonic kidney HEK293T cells (Clontech). HEK293T cells were maintained in T175 flasks (Greiner, 660175) and fed 3 times per week with 30mL of HEK media composed of DMEM/F12 (Thermo Fisher Scientific, 11330-032) with 10% heat inactivated FBS and 1X Glutamax, sodium pyruvate, and non-essential amino acids (Thermo Fisher Scientific, 11140050, 11360070, 35050061). On Day 1 of the protocol, HEK293T cells at 70% confluency were dissociated with 0.25% Trypsin-EDTA (Thermo Fisher, 25200114) for 5 min at room temperature and re-plated at a 1:3 ratio into T175 flasks coated with 0.0001% poly-L-lysine (Sigma, P8920). On Day 2, cells were fed with 20mL HEK media in the morning and transfected in the afternoon using calcium phosphate transfection with viral packaging plasmids (5ug pDMG.2 Addgene#12259, 20ug REV Addgene #12253, and 20ug RRE Addgene#12251) and 20ug of either RTTA or TetO-hNGN2-puro plasmid (Addgene #79049). On Day 3, media was replaced with 30mL fresh HEK media per flask. Viral supernatant was collected and filtered with 0.45μm PVDF syringe filters (EMD Millipore, SLHVR33RS) on days 4 and 5. Lentiviral particles were purified using LentiX Concentrator following manufacturer’s instructions (Takara, 631232). Purified lentiviral particles were resuspended in DMEM/F12 at 10-times the original concentration and stored as single-use aliquots at -80°C. We made 750μL aliquots, which are sufficient to transduce one 6-well plate at 125μL per well.

### Neuronal induction

#### Classic NGN2 Induction

Neurons were generated from iPSCs via direct induction with NGN2. To start the neuronal induction, iPSCs were dissociated with accutase (Stem Cell Technologies, 7920) and seeded on Matrigel-coated 6-well plates at 150,000 cells per well in mTeSR Plus with 10μM Y-27632 (Stem Cell Technologies, 72304). The next day (day 0), each well of iPSCs was transduced with 125μL of tetO-NGN2 and RTTA lentivirus for 2hrs in mTeSR Plus with 10μM Y-27632 at 37°C. After the 2hr incubation, the viral media was aspirated and exchanged for mTeSR Plus with 2ug/mL doxycycline (Stem Cell Technologies, 72742) to start NGN2 expression. On day 1, cells were fed with half mTeSR plus and half Neuronal media, composed of Neurobasal A with 1X B27, NEAA, and Glutamax (Thermo Fisher Scientific, 10888022, 17504044, 35050061, and 11140050) with doxycycline. To select for NGN2-expressing cells, the media was changed on day 2 to neuronal media with doxycycline and 2ug/mL puromycin (Gibco, A11138-03). To minimize batch and cell line variability, we introduced a replating step on day 3 which allowed us to normalize cell density. For this, we coated glass-bottom 96-well plates (Cellvis, P96-1.5H-N) with 50μg/mL poly-D-lysine (Thermo Fisher Scientific, A3890401) in borate buffer (50mM boric acid, 12.5mM sodium tetraborate decahydrate, pH 8.5) for 1hr at 37°C and subsequently coated them with Matrigel. On day 3, induced neurons were re-plated at 2 x 10^4^ cells/well in 96-well plates with neuronal media supplemented with doxycycline and 40nM BRDU (Thermo Fisher Scientific, B23151) to select against the remaining dividing iPSCs. Cells underwent a full media change on days 7 and 10, and a half media change on day 17 with Neuronal media supplemented with 1ug/mL laminin (Sigma, L2020) and 10ng/mL BDNF, GDNF, and NT-3 (PeproTech, 450-02, 450-10, 450-03). Doxycycline was not used past day 7. Neurons were collected on days 23-25. Penicillin-streptomycin was added to all culture media.

#### Enhanced NGN2 Induction

For calcium imaging experiments, we used an extended differentiation protocol inspired by Quist, Ahlenius, and Canals 2021. On Day 7 of the protocol described above, cells underwent a full media change with half Neuronal media and half FGF basal media (Neurobasal, Thermo Fisher, 21103049, with 1% FBS, B27, NEAA, and Glutamax), supplemented with 1μg/mL laminin, 2.5μg/mL doxycycline, 8ng/ml FGF (PeproTech, 100-18B), 5ng/ml CNTF (PeproTech, 450-13), and 10ng/mL each of BDNF, GDNF, NT-3, and BMP4 (Peprotech 120-05ET). From Day 10 onwards, we performed half-media changes every 3-4 days with half Neuronal media and half Maturation media (50% DMEM/F12, 50% Neurobasal, N2, Sodium Pyruvate, Glutamax, with 5μg/ml N-Acetyl-L-cysteine, Sigma-Aldrich, A8199-10G, 5ng/ml Heparin-Binding EGF-like Growth Factor, Sigma-Aldrich, E4643), supplemented with 1μg/mL laminin, 2.5μg/mL doxycycline, 1mM cAMP (Sigma, D0627), and 10ng/ml each of BDNF, GDNF, NT-3, CNTF, and BMP4. Treatments and calcium imaging were performed on days 46-49.

### Microglial induction

iPSC-derived microglial like cells (iMGLs) were generated following the protocol from McQuade and Blurton-Jones 2022 with minor modifications (2). We first generated hematopoietic progenitor cells (HPCs) using the STEMdiff Hematopoietic Kit (Stem Cell Technologies, 05310). In brief, we gently dissociated iPSCs at ∼60-70% confluency into small clusters with 10μM EDTA, seeded 2-4 clusters/cm2 in Matrigel-coated 6-well plates, and performed media changes according to manufacturer’s instructions. From days 3-9, we performed 1mL media additions instead of exchanges. On day 12, floating HPCs were harvested from the media and banked at 2x10^6^ cells/mL in BamBanker freezing media (Bulldog Bio, BB02). A fraction of cells from every batch was seeded on poly-L-lysine-coated plates for immunofluorescent labeling of CD43 as quality control. Immunolabeled HPCs were imaged at 10X Plan Apo Lambda objective (NA = 0.45). To quantify the percentage of CD43-positive cells, we used the Fiji/Image J Cell Counter tool (Version 1.54f), using the signal intensity of samples stained with secondary antibodies only as our background signal threshold.

To generate iMGLs, we seeded 1x10^5^ cells/well in 6-well plates in microglial differentiation media (2).Cells were topped off with 1mL of media per well every other day for 10 days. On day 12, we gently removed 5mL from each well, fed with 1mL of fresh media per well, and continued feeding cells every other day until day 22. On day 24, cells were collected and re-plated in Matrigel-coated 6-well plates at 1x10^6^ cells/well in 1mL of conditioned media plus 1mL of Maturation media. Cells were topped off with 1mL of Maturation media every other day until day 30. iMGLs were collected on day 31. Penicillin-streptomycin was added to all culture media throughout the differentiation. For immunocytochemistry, iMGLs were re-plated on poly-D-lysine coated 96-well plates, allowed to attach in the incubator for at least 4hrs, and fixed for immunolabeling.

### iMGL stimulation with LPS

On day 24 of microglia differentiation, iMGLs were treated with either 50ng/mL lipopolysaccharide (LPS; Sigma, L4516) or vehicle (PBS). Treatment was administered with every feeding for a week on days 24, 26, 28, and 30.

### iMGL phagocytosis assay

Fluorescent latex beads (Sigma, L2778) were pre-opsonized in FBS at a 1:6 ratio for 30min. Beads were then diluted with microglia Maturation media to get a stock concentration of 0.01%. Cells were treated at a final concentration of 0.0003% beads and incubated for 16hrs at 37°C. Cells were harvested and kept on ice until fixed with 4% paraformaldehyde (Electron Microscopy Sciences, 50980487). For negative controls, iMGLs were treated with beads, immediately placed on ice, and fixed. Cells were washed and resuspended in Sorting Buffer (2.5mM EDTA, 25mM HEPEs pH 7.0, 1% penicillin/streptomycin in 1X PBS). Phagocytic cells were quantified by flow cytometry (Bio-Rad ZE5 Cell Analyzer).

### Reverse transcription qPCR

We extracted mRNA from cells using a kit and following manufacturer’s instructions (Zymo Research, NC1000412). We performed reverse transcription using the SuperScript IV First Strand Synthesis System (Thermo Fisher Scientific, 18091050) and made single-use cDNA aliquots stored at 20°C. Quantitative reverse transcription PCRs (RT-qPCRs) were done with 2ng of cDNA per reaction using SYBR Green Universal Master mix (Bio-Rad, 1725122) and a CFX384 Real Time PCR System (Bio-Rad). Primers listed in SI Appendix Table S1.

### Calcium imaging with Fluo-4AM

For calcium imaging, induced neurons were grown in 96-well plates with #1.5 cover glass bottoms (Cellvis, NC0536760). Fluo4-AM was used according to manufacturer’s instructions (Thermo Fisher Scientific, F14201). In brief, Fluo4-AM was reconstituted with 4.56μL of Pluronic F127 20% solution in DMSO (Thermo Fisher Scientific, P3000MP) and 4.56μL of DMSO (Thermo Fisher Scientific, BP231100). This stock solution was diluted 1:1000 in Imaging Buffer (130mM NaCl, 4mM KCl, 2mM CaCl2, 1mM MgCl2, 10mM HEPES, and 10mM glucose). Induced neurons were incubated in Fluo4-AM working solution for 30min at 37°C, washed twice with Imaging Buffer, and incubated again at 37°C for 20min to allow for the de-esterification of the calcium indicator. Time-lapse imaging was done in an Image Xpress Confocal HT.ai high-content imaging system (Molecular Devices) with CO_2_ incubation at 37°C. We used a 10X Plan Apo Lambda objective (NA = 0.45) to capture 2-minute recordings from 2-3 fields of view per well at 20 frames per second. Cells were imaged with the FITC laser at 1% power to minimize photobleaching.

### Calcium imaging data processing

Time-lapse recordings were concatenated and trimmed to 2000 frames (equivalent to 100 sec) using Fiji/Image J Version 1.54f. We used the analysis pipeline Suite2p (https://github.com/MouseLand/suite2p) to identify cell regions of interest (ROIs) and extract fluorescent signals. Further analysis was performed with Python 3 and Microsoft Excel. To quantify calcium signals, we calculated ΔF/F_0_ = (F_t_-F_0_)/F_0_, where F_t_ is the fluorescent signal at time t and F_0_ is the average fluorescence over a baseline period of 0.5s (10 frames), selected as the 0.5s window with the lowest fluorescent signal for each ROI. Normalized calcium signals were smoothened with a rolling average of 10 frames and calcium spikes were identified with a rolling maximum of 100 frames. To filter out noise, we set a minimum amplitude threshold equivalent to 2 standard deviations from the mean baseline signal. Based on the premise that neuronal firing is accompanied by a rapid change in fluorescent signal, we identified the timing of spike initiation by calculating the point at which ΔF first exceeded 5 standard deviations from the baseline ΔF in a rolling window of 50 frames. We measured network activity by quantifying two types of correlated activity: coordinated and synchronized events. We defined coordinated events as instances where >50% of cell ROIs in a field of view (FOV) had a spike initiation within a 250msec window (5 frames) and synchronized events as instances where >50% of cell ROIs in a FOV had a spike initiation within a narrower window of 50msec (1 frame).

### Immunofluorescent labeling

For all immunofluorescence assays, cells were plated on glass-bottom 96-well plates (Cellvis, NC0536760). Cells were fixed with 4% PFA in PBS for 20 minutes and washed once with PBS. For immunolabeling of cytosolic proteins, cells were blocked and permeabilized in PBS with 3% bovine serum albumin (BSA, Sigma, A7906), and 0.01% TritonX-100 (Sigma, T9284) for 30 minutes at room temperature. For labeling surface markers, blocking was done without detergent. After blocking, cells were incubated with primary antibody in 3% BSA overnight at 4°C, washed twice with PBS, and incubated with secondary antibody and/or Bodipy-493/503 (1μg/mL, Invitrogen, D3922) in PBS for 1hr. To label nuclei, cells were incubated with DAPI (1:1000, Invitrogen, D1306) for 5-10 minutes. Cells were then washed once more with PBS before imaging. Primary antibodies used in this study: Nanog (1:500, Thermo Fisher, PA5-85110), Oct4 (1:100, Thermo Fisher, MA1-104), Sox2 (1:400, Cell Signaling, 3579), and SSEA4 (1:150, Abcam, 16287), βIII-tubulin (1:250, Synaptic Systems, 302304), NeuN (1:500, Sigma, MAB377), Cult1 (1:500, Abnova, H00001523-M01), vGlut1 (1:500, Synaptic Systems, 135318), vGlut2 (1:500, Novus Biologicals, NBP2-94571), Gad67 (1:500, EMD Millipore, MAB5406MI), synapsin 1/2 (1:1000, Synaptic Systems, 106002), CD43 (1:100, R&D Systems, MAB2038), PU.1 (1:100, Cell Signaling Technologies, 2266), Iba1 (1:100, Fisher Scientific, PIPA518488), P2RY12 (1:50, Sigma, HPA014518), Trem2 (1:100, R&D Systems, AF1828). For secondary antibodies, we used the Alexa fluor conjugated antibodies (1:1000, Life Technologies, A21202, A11008, A11036, A11031, A21450, A31571, A31573).

### High content confocal imaging

Confocal imaging was done using an Image Xpress Confocal HT.ai high-content imaging system (Molecular Devices). We used the following objectives: 10X (NA = 0.45) Air Plan Apo Lambda, 20X (NA = 0.95) and 40X (NA = 1.15) water immersion Apo Lambda S LWD objectives.

### Image Analysis

All image analysis was performed using IN Carta Image Analysis Software (v2.5.09062223, Molecular Devices).

#### Iba1 and PU.1

To quantify the percentage of iMGLs expressing PU.1 and Iba1, we took z-stacks with 5 steps at 1μm intervals using a 20X objective. We generated maximum intensity projections and used the IN Carta Nuclei.a.h5 SINAP model to identify nuclei ROIs based on DAPI stain. We then measured the signal intensity of the PU.1 and Iba1 channel inside the nuclei ROIs. We used the DT-Classifier tool to set fluorescent signal intensity thresholds for categorizing PU.1 positive and negative cells, and for sub-dividing PU.1 positive ROIs into Iba1 positive and negative ROIs. We used secondary only controls to find the average background signal and used that to set the thresholds.

#### NeuN and Cux1

To quantify the percentage of iNs expressing NeuN and Cux1, we took z-stacks with 10 steps at 0.5μm intervals using a 20X objective. We then generated maximum intensity projections and used the IN Carta Nuclei.a.h5 SINAP model to identify nuclei ROIs based on DAPI stain. We then measured the signal intensity of the NeuN and Cux1 channels inside the nuclei ROIs. We used the DT-Classifier tool to set thresholds and categorize NeuN- and Cux1-positive cells.

#### Lipid droplets

We used two different methods for the quantification of lipid droplets (LDs). To obtain LDs normalized to cell count, we took a stack of 10 sequential z-steps every 0.5μm at 20X and generated maximum intensity projections. LDs were segmented using the IN Carta Robust Puncta algorithm, with a size threshold to avoid detecting large, non-LD structures. Healthy nuclei were segmented with a modified version of the IN Carta Nuclei.a.h5 SINAP model, re-trained with our own data to better identify healthy nuclei and avoid condensed nuclei and debris. To obtain LDs normalized to neurite area, iNs were imaged at 40X on a single z-plane. LDs were segmented as described above and βIII-tubulin immunolabeling was segmented using the IN Carta Robust algorithm for neurites.

#### Synaptic puncta

To quantify synaptic puncta, we took single-plane images at 40X of iNs immunolabeled for synapsin 1/2 and βIII-tubulin and stained with DAPI. Synapsin puncta were segmented using the IN Carta Robust Puncta algorithm, with thresholds for signal intensity and particle size to avoid detection of larger structures.

### Isolation of exosomes

Exosomes were isolated from microglia conditioned media using sequential ultracentrifugation. Conditioned media was harvested from iMGL cultures in 50mL conical tubes, centrifuged at 300 x g for 5 minutes to remove cells and then at 2,000 x g on a benchtop centrifuge (Eppendorf) to remove debris. 36mL of conditioned media per condition was then transferred into a polypropylene centrifuge tube (Beckman Coulter, 326823) and centrifuged at 10,000 x g for 45 minutes using an Optima XE-100 centrifuge with an SW32-Ti swinging bucket rotor (Beckman Coulter). This step yields a pellet of microvesicles, with exosomes remaining in the supernatant. The supernatant was carefully transferred to an ultra-clear centrifuge tube (Beckman Coulter, 344058) and centrifuged at 100,000 x g for 1.5hrs to pellet exosomes. The supernatant was carefully aspirated and discarded, and the exosome pellet was washed with 35mL of cold PBS with Ca^2+^ Mg^2+^ (Life Technologies, 14040117) and centrifuged at 100,000 x g for 1hr. This step was repeated, and the resulting exosome pellet was resuspended in 50μL PBS with Ca^2+^ Mg^2+^. Conditioned media and exosome samples were kept on ice throughout the purification process and all centrifugation steps were performed at 4°C.

### Exosome treatments

For exosome treatments, 15μL of resuspended exosomes were used to perform a protein assay (DC Protein Assay, Bio-Rad, 5000112). Exosome samples were then diluted with PBS with Ca^2+^ Mg^2+^ to achieve a concentration of 0.68μg/mL. Each well of iNs (150μL neuronal media) was treated with 5μL of EVs.

### Exosome Western blots

To characterize microglial small EVs, we isolated EVs from 35mL of conditioned media from D24 iMGLs and resuspended them in 50μL RIPA buffer with protease inhibitors (Thermo Fisher Scientific, A32955). Samples were sonicated for 10 pulses using a wand sonic dismembrator (Fisher Scientific Model 100, setting 3). 35μL of each sample was mixed with 6X Laemmli buffer (Thermo Fisher Scientific, J61337AC) before boiling for 10min and loading into a 4-20% Tris-glycine gel (Bio-Rad, 4561094). We ran the gel at 100V for 2hrs then transferred using a Trans-Blot Turbo Transfer system with a mini transfer pack (Bio-Rad, 1704158). We incubated the membrane in Ponceau S solution for 1-5min to visualize total protein. The membrane was cut between the 75 and 100kDa markers and the sections were blocked separately in blocking buffer (5% milk in TBS with 0.05% Tween20—Fisher, BP337-100) for 1hr at room temperature. Membranes were then incubated in fresh blocking buffer with primary antibodies against Alix (1:1000, EMD Millipore, ABC40) and Flotillin-1 (1:1000, Sigma, F1180) overnight at 4°C. The next day, the membranes were washed 3 x 10min in TBS with 0.05% Tween20 and incubated in blocking buffer with HRP-conjugated secondary antibody (1:1000, Bio-Rad, 1706515) for 1hr at room temperature. After another 3 washes with TBS-tween20, membranes were developed using a Super Signal West Femto kit (Thermo Fisher, 34095) according to manufacturer’s instructions.

### Negative stain transmission electron microscopy (TEM)

Carbon-coated copper grids (400 mesh) were glow-discharged and 10 µL of each sample (purified EVs resuspended PBS with Ca^2+^ Mg^2+^) was adsorbed for 2 minutes. Excess sample was wicked away and grids were negatively stained with 2% uranyl acetate for 2 minutes. Excess stain was wicked away and the grids were allowed to dry. Samples were analyzed at 80kV with a Thermo Fisher Talos L120C transmission electron microscope and images were acquired with a CETA 16M CMOS camera.

### Statistical Analysis

All statistical analyses were done using GraphPad Prism software version 10.4.1 (GraphPad Software Inc.). Statistical significance was assessed using one-way or two-way ANOVA. One-way ANOVA was used for comparisons of 3 or more groups, followed by Tukey or Sidak post hoc tests. Two-way ANOVA was used for comparisons with 2 variables and was followed by Tukey post hoc tests. Data are represented as mean ± SEM. A p-value < 0.05 was considered significantly different and p-value was noted as follows: * <0.05, ** <0.01, *** <0.001, and **** <0.0001.

---

**Fig. S1.**
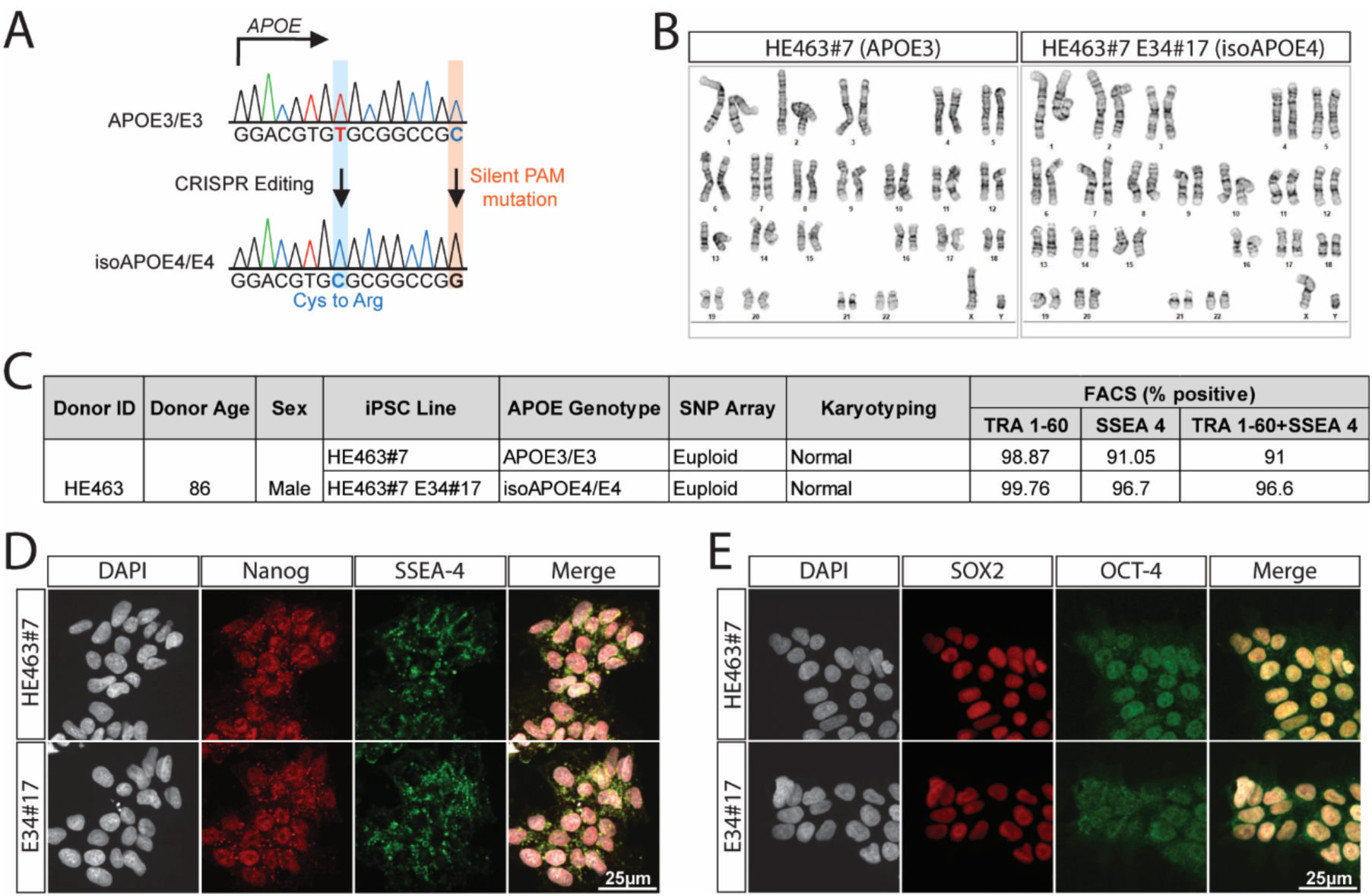
Characterization of iPSC lines. A) We used an iPSC line with APOE3/E3 genotype (HE463#7) and CRISPR-edited it to change amino acid residue 112 from a cysteine to an arginine and generate an isogenic APOE4/E4 iPSC line (HE463#7 E34#17). We also introduced a non-coding mutation at the protospacer adjacent motif (PAM) site to prevent recurrent Cas9 cutting. B) Karyographs for both HE463#7 and HE463#7 E34#17 iPSC lines, showing normal karyotypes. C) Table showing donor information and characterization of iPSC lines. Both lines were screened for chromosomal abnormalities via SNP array and for pluripotency markers TRA 1-60 and SSEA-4 via flow cytometry. D) Cropped confocal images of iPSCs stained with DAPI and immunolabeled for Nanog and SSEA-4, and E) Sox2 and OCT-4.

**Fig. S2.**
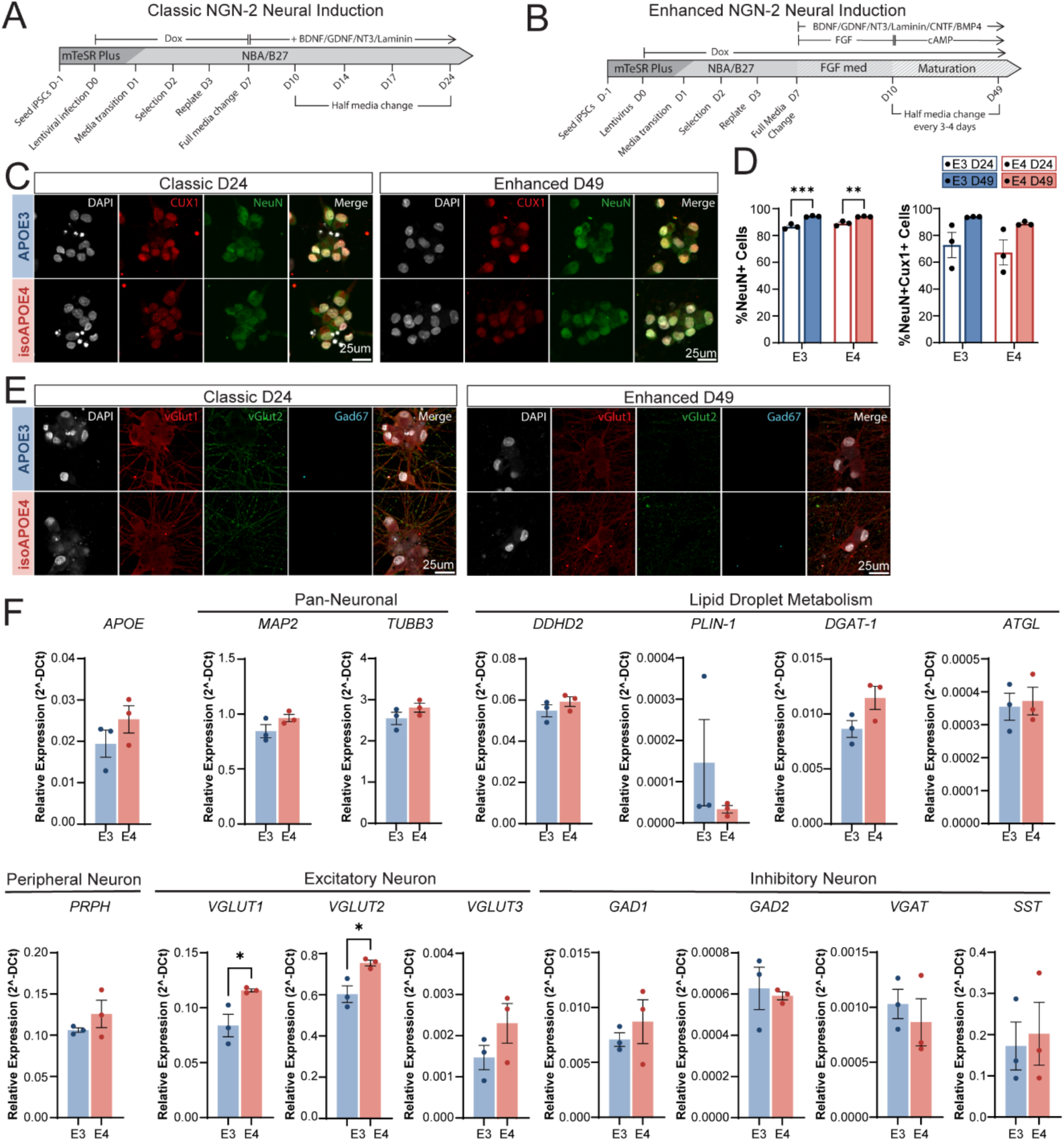
Characterization of iNs. iNs were generated from APOE3 (HE463#7) and isoAPOE4 (HE463#7 E34#17) iPSCs. A) Differentiation timeline for Classic and B) Enhanced NGN-2 neural inductions with modifications inspired by Canals et al. 2018. C) Cropped representative confocal images of immunofluorescent labeling of iNs generated with Classic (left) and Enhanced (right) induction protocols. iNs were labeled for Cux1 (red), NeuN (green), and nuclei were stained with DAPI (white). D) Quantification of NeuN+ (top) and NeuN+CUX1+ (bottom) cells generated with both protocols. Data points represent well averages (N = 3-4 culture wells per condition; Two-way ANOVA with Sidak correction for multiple comparisons.). E) Immunocytochemistry for excitatory synaptic markers vGlut1 (red) and vGlut2 (green) and inhibitory synapse marker Gad67 (cyan). Confocal images are cropped for visualization. F) RT-qPCR characterization of iNs generated with Classic NGN-2 neural induction protocol at D24. Plots show relative expression (2^ΔCt) normalized to housekeeping gene *PPIA* (N = 3 independent experiments). Unpaired T-test. All bar graphs represent the mean with error bars representing SEM. * p < 0.05, ** p < 0.01, and *** p < 0.001.

**Fig. S3.**
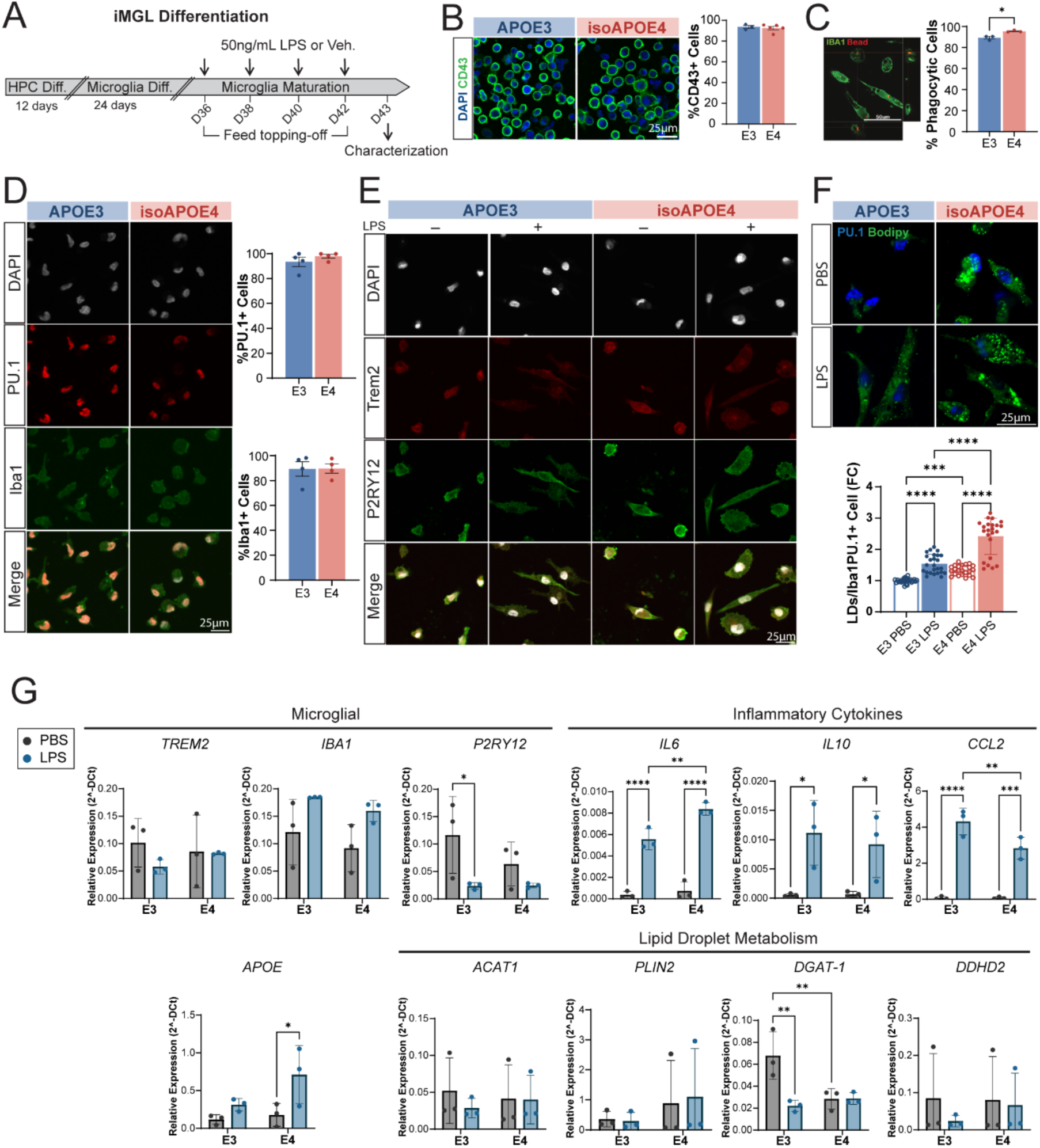
Characterization of iMGLs and LPS-stimulation paradigm. A) Timeline of microglial differentiation. iMGLs were generated from iPSCs following the protocol from McQuade et al., 2018. On days 36, 38, 40, and 42, iMGLs received 50ng/mL LPS or vehicle (PBS) at the time of feeding, and they were characterized on D43. B) HPCs at day 12 of differentiation from iPSCs were fluorescently labeled for CD43 (green) and DAPI (blue) and CD43+ cells were quantified (right; N = 3-5 independent preps, unpaired t test). C) Confocal image of iMGLs immunolabeled for Iba1 with phagocytosed fluorescent latex beads (left) and quantification of the percentage of phagocytic cells obtained via flow cytometry. (right; N = 3 independent preps, Unpaired t test). D) Representative confocal images of iMGLs immunolabeled for PU.1 (red) and Iba1 (green) and stained with DAPI (white), with quantification of the percentage of PU.1- and Iba1-expressing iMGLs (right; N = 4 independent preps; Unpaired t test). E) Immunofluorescent labeling of Trem2 (red) and P2RY12 (green) in iMGLs with a 7-day treatment of either PBS or 50ng/mL LPS. F) Representative images of iMGLs with immunolabeling of PU.1 (blue) and BODIPY-493/503 labeling of LDs. LDs were quantified per cell and plotted as the fold-change relative to the average of the E3 PBS condition (N = 23-25 wells from 4 independent experiments; Two-way ANOVA with Fisher’s LSD Test). G) RT-qPCR characterization for D31 iMGLs treated with PBS or LPS. Plots show relative expression (2^ΔCt) to housekeeping gene PP1A (N = 3 independent experiments; Two-way ANOVA with Fisher’s LSD Test). All bar graphs represent the mean with error bars representing SEM. * p < 0.05, ** p < 0.01, *** p < 0.001, and **** p < 0.0001.

**Fig. S4.**
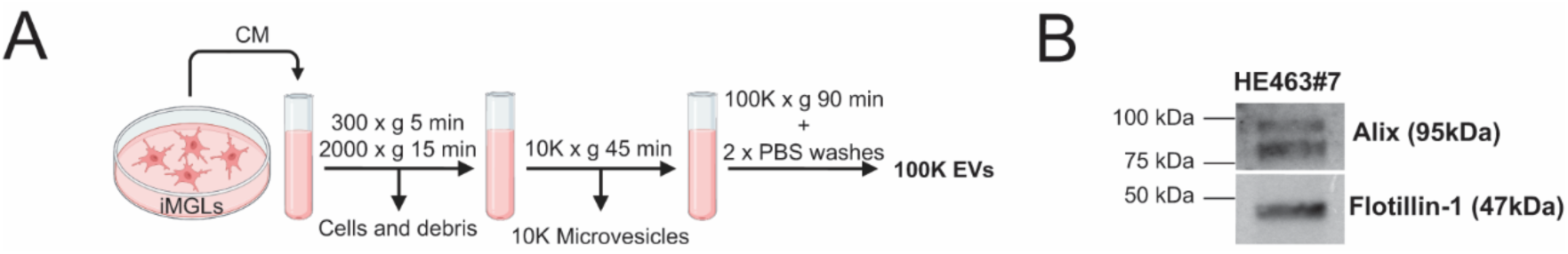
Exosome isolation and characterization. A) Strategy for isolating exosomes from iMGL conditioned media via sequential ultracentrifugation. B) Western blot of purified exosomes showing expression of Alix and Flotillin-1. Alix appears as two bands corresponding to its opened and closed conformations (3).

**Fig. S5.**
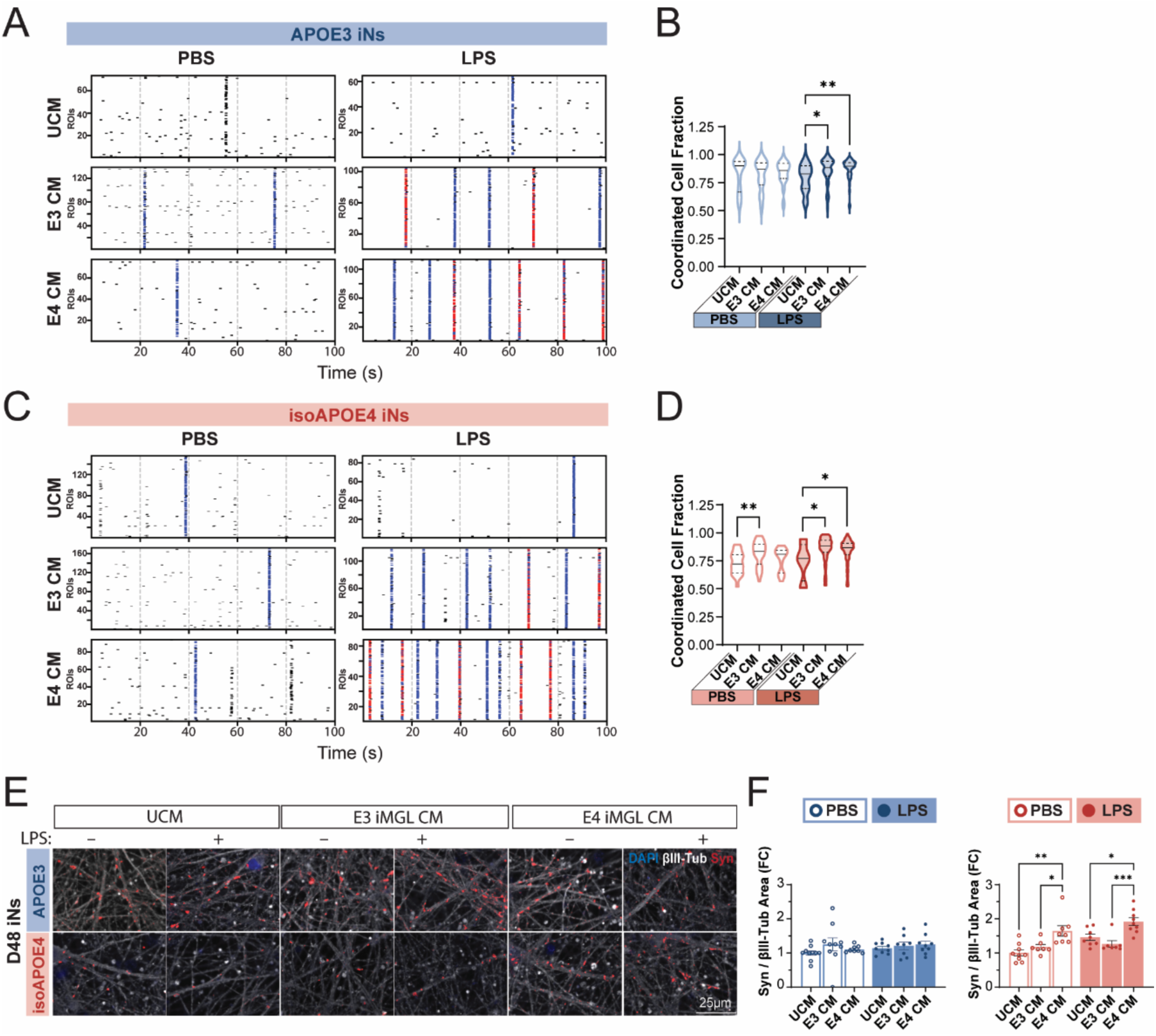
Microglial CM signaling recruits neurons into co-active networks and increases pre-synaptic puncta density in inflammatory state- and APOE genotype-dependent way. A,C) Representative raster plots of Ca-imaging recordings from APOE3 (A) and isogenic APOE4 (C) neurons treated with UCM or CM from PBS- or LPS-treated iMGLs, shown in Fig. 2. Blue tick marks represent coordinated events, defined as instances where >50% of cells in the field of view (FOV) spiked within a 250ms window (5 frames). Red tick marks represent synchronized events, instances where >50% of cells spiked within a 50ms window (1 frame). B,D) Violin plots showing the fraction of cells engaged in each coordinated event. Median, 25%, and 75% quartile values for APOE3 iNs (B): PBS UCM, 0.90, 0.67, 0.94; E3 PBS CM, 0.87, 0.73, 0.93; E4 PBS CM, 0.86, 0.79, 0.92; LPS UCM, 0.83, 0.70, 0.90; E3 LPS CM, 0.88, 0.81, 0.94; E4 LPS CM, 0.90, 0.84, 0.93; isoAPOE4 iNs (D): PBS UCM, 0.72, 0.64, 0.80; E3 PBS CM, 0.83, 0.72, 0.90; E4 PBS CM, 0.81, 0. 64, 0.84; LPS UCM, 0.77, 0.57, 0.89; E3 LPS CM, 0.88, 0.83, 0.93; E4 LPS CM, 0.87, 0.81, 0.91. (N = 14-227 coordinated events from 4 independent experiments). E) Representative confocal images of iNs immunolabeled for synapsin-1/2 (red), βIII-tubulin (white), and stained with DAPI (blue). Images were taken at 40X magnification and cropped for visualization. F) Quantification of synapsin puncta per unit area of βIII-tubulin for APOE3 (left) and isogenic APOE4 (right) iNs (cell lines HE463#7 and E34#17). Data was normalized to the average of the UCM PBS condition and is shown as fold-change (N = 7-10 wells from 2 independent experiments). Statistical comparisons were performed separately within the PBS-treated and LPS-treated groups using one-way ANOVAs followed by Tukey’s post hoc tests. All bar graphs represent mean with error bars representing SEM. * p < 0.05, ** p < 0.01, *** p < 0.001, and **** p < 0.0001.

**Fig. S6.**
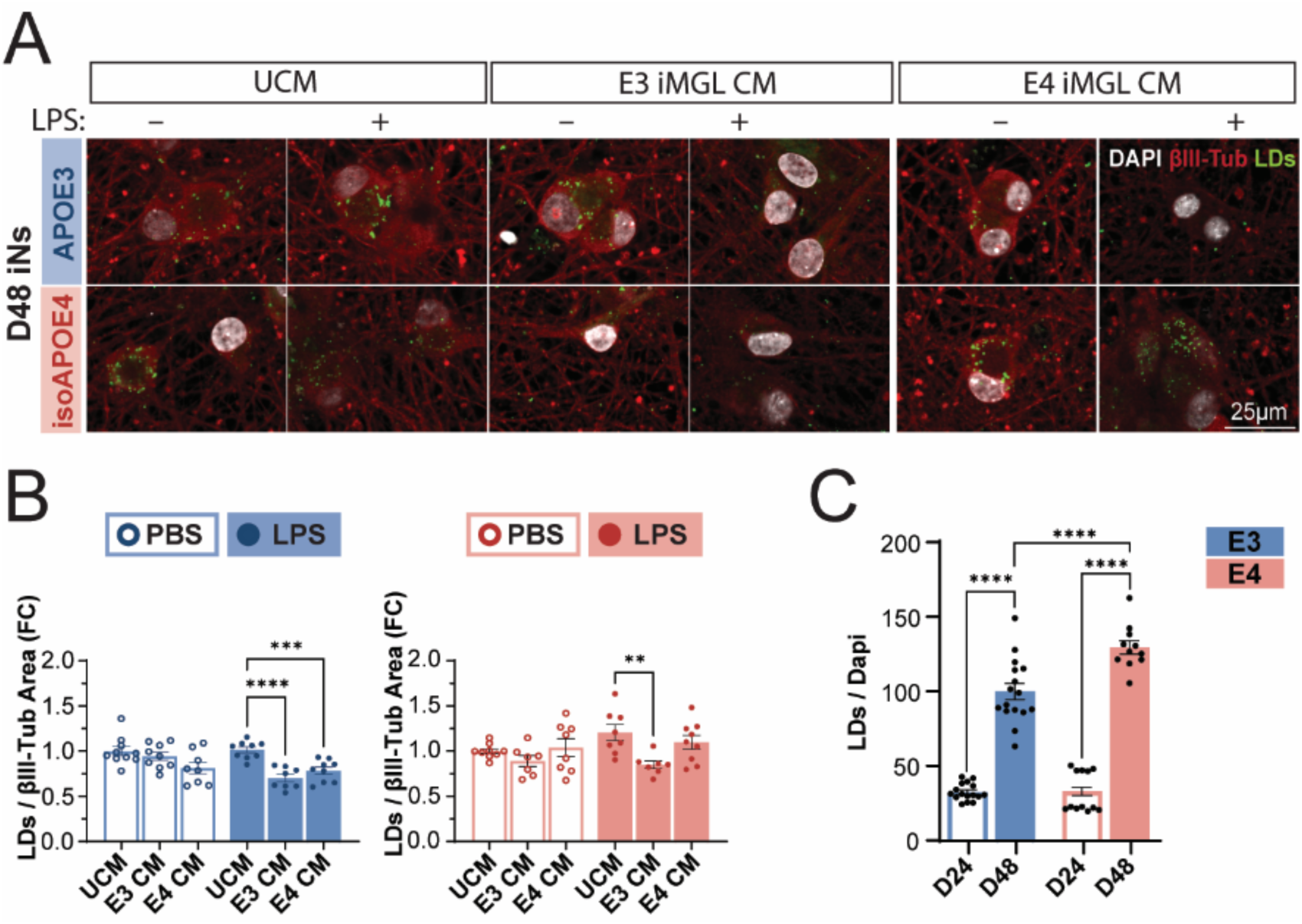
Inflammatory iMGL conditioned media decreases neuronal lipid droplet load, analyzed as neuronal lipid droplets normalized to βIII-tubulin. A) Representative confocal images of D48 APOE3 and iso APOE4 iNs immunolabeled for βIII-tubulin (red) and labeled with DAPI (nuclei, white) and BODIPY-493/503 (LDs, green). Cultures were treated with UCM, E3 iMGL ± LPS CM or E4 iMGL ± LPS CM as indicated. Images were collected at 40X magnification and cropped for visualization. B) Quantification of BODIPY-labeled lipid droplets (LDs) normalized to unit area of βIII-tubulin for APOE3 (left, blue) and isogenic APOE4 (right, red) iNs. This is an alternate analysis method to that of Figure 3. Data points represent well averages and are shown as the fold-change relative to the average of the UCM PBS condition (N = 7-10 wells from 2 independent experiments; Statistical comparisons were performed separately within the PBS-treated and LPS-treated groups using one-way ANOVAs followed by Tukey’s post hoc tests). C) D24 iNs have significantly lower LD load than D48 iNs. Quantification of average iN LDs per well normalized to DAPI-labeled nuclei in D24 and D48 cultures (N = 11-17 wells from 2-3 independent experiments; Two-way ANOVA with Sidak correction for multiple comparisons). All bar graphs represent mean with error bars representing SEM. ** p < 0.01, *** p < 0.001, and **** p < 0.0001.

**Fig. S7.**
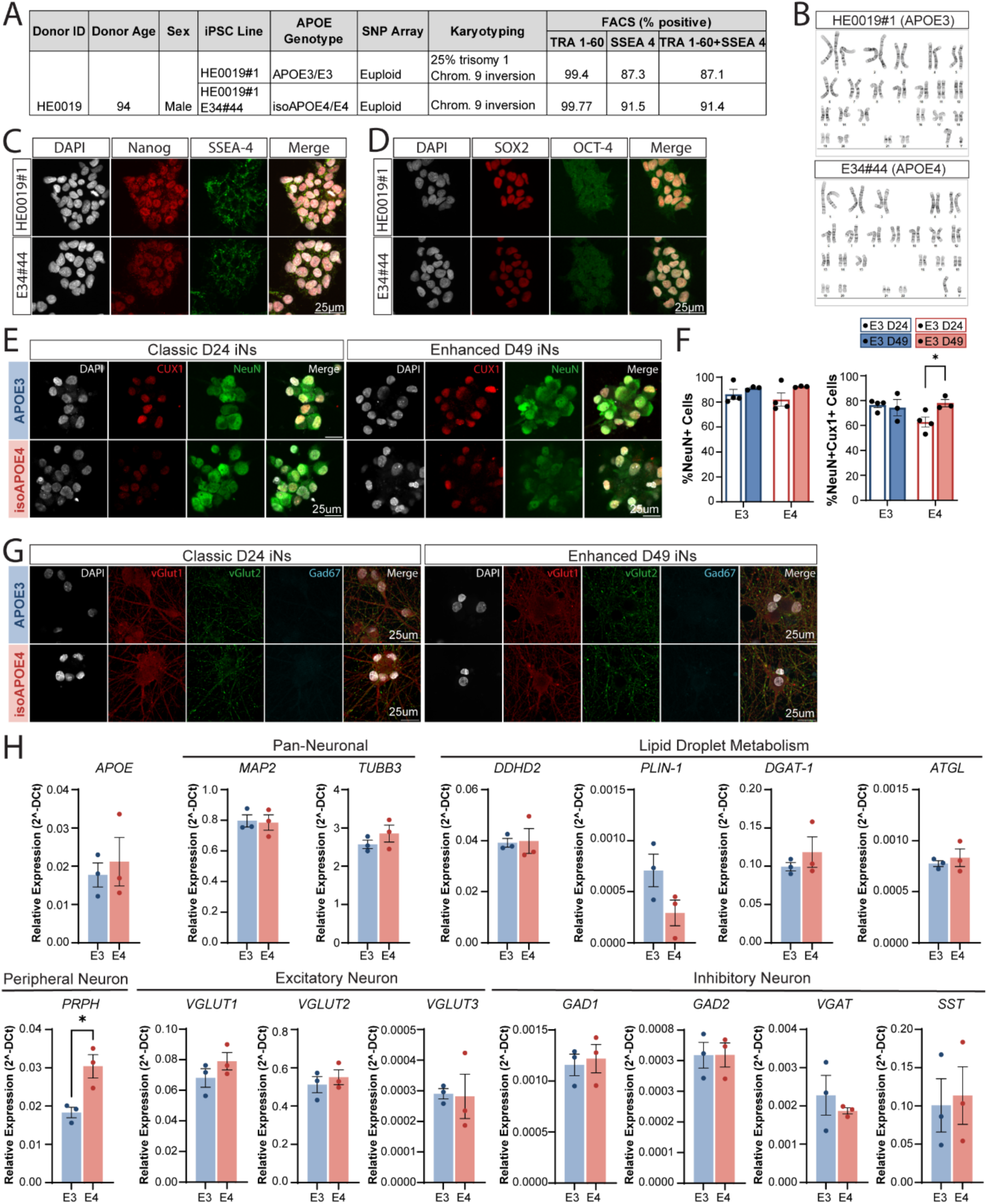
Characterization of additional iPSC lines and iNs. A) Table with donor information and characterization for additional APOE3/E3 iPSC line (HE0019#1) and its isogenic APOE4/E4 iPSC line (HE0019#1 E34#44). Both lines were screened for chromosomal abnormalities via SNP array and karyotyped. Karyotyping revealed 25% of HE0019#1 cells had trisomy 1. Both lines have a pericentric inversion on chromosome 9, which is a common population variant. iPSC lines were also screened for pluripotency markers TRA 1-60 and SSEA-4 via flow cytometry. B) Karyographs for both HE0019#1 and HE0019#1 E34#44 iPSC lines. C) Cropped confocal images of iPSCs stained with DAPI and immunolabeled for Nanog and SSEA-4, and D) Sox2 and OCT-4. E) Cropped representative confocal images of immunofluorescent labeling of iNs generated with Classic (left) and Enhanced (right) induction protocols. iNs were labeled for Cux1 (red), NeuN (green), and nuclei were stained with DAPI (white). F) Quantification of NeuN+ (left) and NeuN+CUX1+ (right) cells generated with both protocols. Data points represent well averages (N = 3-4 culture wells per condition; Two-way ANOVA with Sidak correction for multiple comparisons.). G) Immunocytochemistry for excitatory synaptic markers vGlut1 (red) and vGlut2 (green) and inhibitory synapse marker Gad67 (cyan). Confocal images are cropped for visualization. H) RT-qPCR characterization of iNs generated with Classic NGN-2 neural induction protocol at D24. Plots show relative expression (2^ΔCt) normalized to housekeeping gene *PPIA* (N = 3 independent experiments; Unpaired T-test). All bar graphs represent the mean with error bars representing SEM. * p < 0.05.

**Fig. S8.**
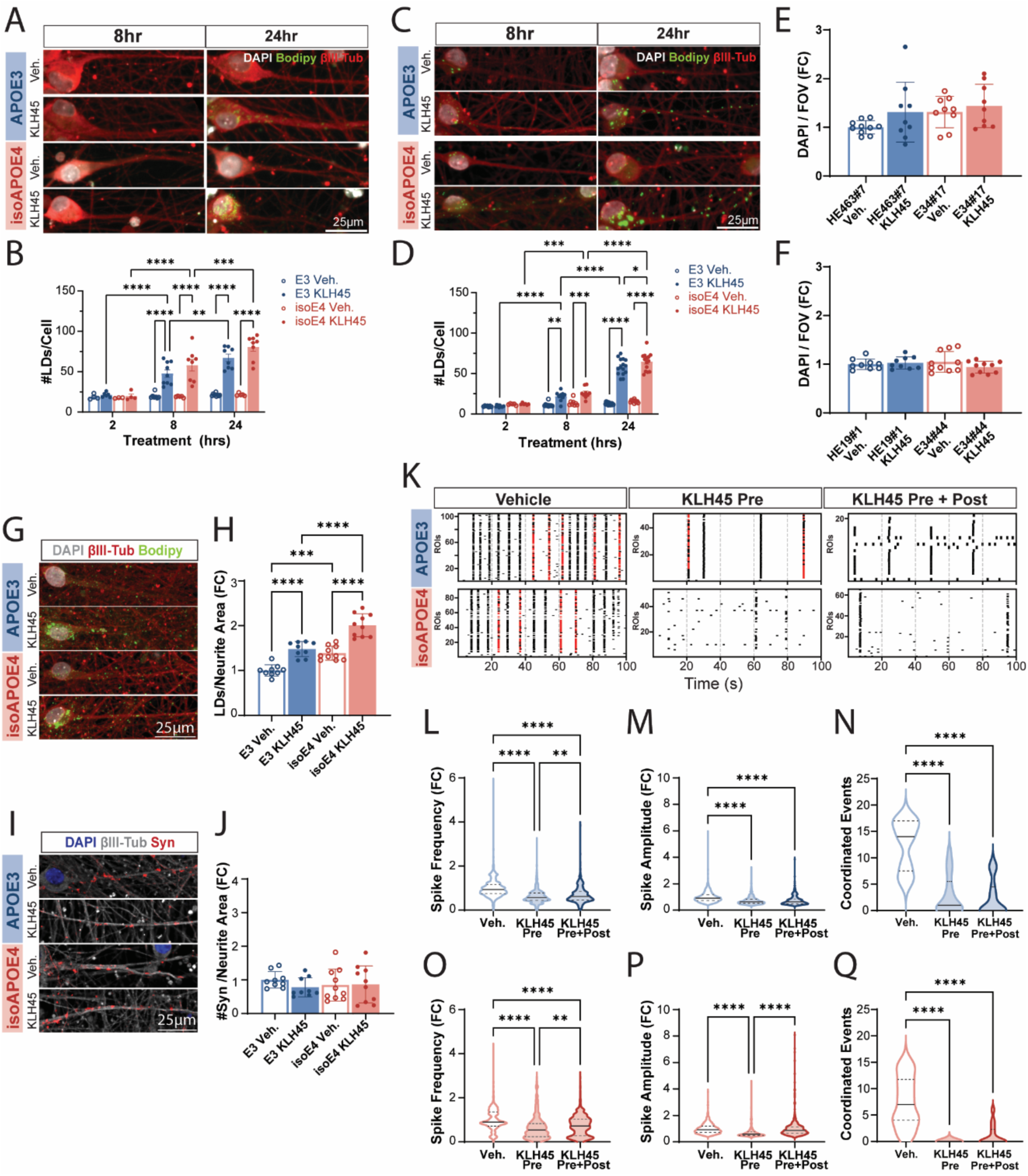
Short KLH45 treatment timepoints and validation of KLH45 treatments in additional iN lines. A-D) D23 iNs generated with the Classic NGN-2 neural induction protocol from HE463#7/E34#17 (A-B) and HE19#1/E34#44 (C-D) lines were treated with 5μM KLH45 or vehicle (EtOH) for 2, 8, or 24hrs. A/C) Cropped representative images of iNs labeled with DAPI (white), Bodipy-488 (green), and immunolabeled for βIII-tubulin (red). Images are max intensity projections of 10μm confocal z-stacks taken at 1μm steps and cropped for visualization. B/D) Quantification of lipid droplets per field of view normalized to DAPI count and averaged by well. Analyzed fields of view were 345μmx345μm (B: N = 3-9 wells from 2 independent experiments; D: N = 5-15 wells from 3 independent experiments; Two-way ANOVA with Sidak correction for multiple comparisons). E-F) Quantification of healthy nuclei based on DAPI stain per field of view for HE463#7/E34#17 (E; N = 9-10 wells from 2 independent experiments; Two-way ANOVA with Sidak correction for multiple comparisons) and HE19#1/E34#44 iNs (F; N = 9-10 wells from 2 independent experiments; Two-way ANOVA with Sidak correction for multiple comparisons) after a 24hr treatment with either vehicle (EtOH) or 5μM KLH45. G-Q) Validation of experiments in Figure 4 with independent iPSC lines HE0019#1 (APOE3) and E34#44 (isoAPOE4). D49 iNs were generated with the Enhanced NGN-2 induction protocol. G) Representative confocal images of iNs treated with vehicle or KLH45 for 24hrs, labeled with DAPI (white), Bodipy-488 (green), and immunolabeled for βIII-tubulin (red). Images were cropped for visualization. H) Quantification of lipid droplets per field of view, normalized to total neurite area and shown as the fold change relative to the average of E3 Veh-treated iNs (N = 9-10 wells from 2 independent experiments; Two-way ANOVA with Sidak correction for multiple comparisons). I) Representative images of D49 iNs stained with DAPI (blue) and immunolabeled for βIII-tubulin (white) and synapsin 1/2 (red). Images are max intensity projections of 10μm confocal z-stacks taken at 1μm steps and cropped for visualization. J) Quantification of synapsin puncta per field of view, normalized to total neurite area and shown as the fold change relative to the average of E3 Veh-treated iNs. (N = 9-10 wells from 2 independent experiments; Two-way ANOVA with Sidak correction for multiple comparisons). K) Representative raster plots of Ca-imaging with Fluo4-AM in D49 iNs treated as described in Figure 4. Red tick marks represent synchronized events, which we defined as events as instances where >40% of neurons in the field of view spiked within a 50ms window. L-Q) Quantification of Ca-imaging features for APOE3 (L-N; blue) and isoAPOE4 (O-Q; red). L/O) Violin plots showing average spike frequency per cell normalized as fold change to the average of Veh-treated iNs. Median, 25%, and 75% quartile values for APOE3 iNs (L): Veh, 0.94, 0.76, 1.17; KLH45 Pre, 0.57, 0.45, 0.78; KLH45 Pre+Post, 0.62, 0.47, 0.86; isoAPOE4 iNs (O): Veh, 0.90, 0.72, 1.36; KLH45 Pre, 0.54, 0.24, 0.83; KLH45 Pre+Post, 0.72, 0.27, 1.04 (N = 505-1877 cells from 2 independent experiments). M/P) Violin plots showing average spike amplitudes per cell normalized as fold change to the average of Veh-treated iNs. Median, 25%, and 75% quartile values for APOE3 iNs (M): Veh, 0.91, 0.73, 1.17; KLH45 Pre, 0.64, 0.52, 0.83; KLH45 Pre+Post, 0.64, 0.49, 0.85; isoAPOE4 iNs (P): Veh, 0.92, 0.71, 1.19; KLH45 Pre, 0.58, 0.50, 0.75; KLH45 Pre+Post, 0.86, 0.65, 1.14 (N = 505-1871 cells from 2 independent experiments). N/Q) Violin plots showing quantification of coordinated events per 100sec recording session. Median, 25%, and 75% quartile values for APOE3 iNs (N): Veh, 14, 7.5, 17; KLH45 Pre, 1, 0, 5.5; KLH45 Pre+Post, 0, 0, 4.5; isoAPOE4 iNs (Q): Veh, 7, 4, 11.75; KLH45 Pre, 0, 0, 0; KLH45 Pre+Post, 0, 0, 2.25 (N = 14-16 recordings, 2 recordings per well from 2 independent experiments). One-way ANOVA with Tukey correction for multiple comparisons. Bar graphs represent mean with SEM. Violin plot lines represent median (solid) and quartiles (dashed). * p < 0.05, ** p < 0.01, *** p < 0.001, and **** p < 0.0001.

**Table S1.**
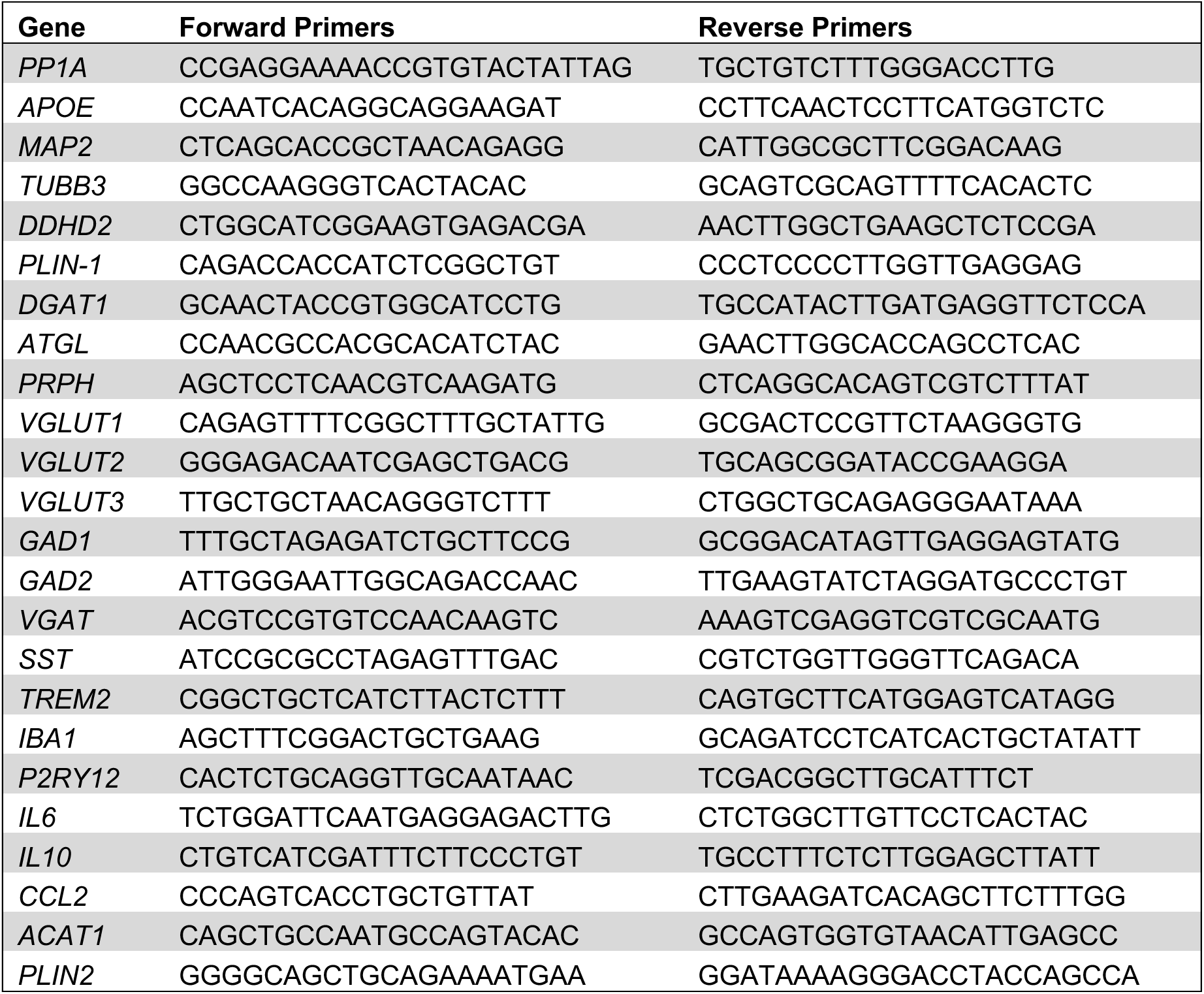
Primers for RT-qPCR.

**Table S2.** Antibodies and Dyes.

| **Antibody/ Dye** | **Company** | **Catalog #** | **Dilution** |
| --- | --- | --- | --- |
| DAPI | Invitrogen | D1306 | 1:1000 |
| Fluo-4AM | Thermo Fisher Scientific | F14201 | N/A |
| Alexa Fluor 488 anti-human TRA-1-60-R | BioLegend | 330613 | 1:20 |
| APC anti-human SSEA-4 | BioLegend | 330417 | 1:20 |
| rabbit anti-Nanog | Thermo Fisher Scientific | PA5-85110 | 1:500 |
| mouse anti-Oct4 (9B7) | Thermo Fisher Scientific | MA1-104 | 1:100 |
| rabbit anti-Sox2 | Cell Signaling | 3579 | 1:400 |
| mouse anti-SSEA4 | Abcam | 16287 | 1:150 |
| guinea pig anti-BIII Tubulin | Synaptic Systems | 302 304 | 1:250 |
| mouse anti-NeuN | Sigma-Aldrich | MAB377 | 1:500 |
| rabbit anti-Cult1 | Abnova | H00001523-M01 | 1:500 |
| guinea pig anti-vGlut1 | Synaptic Systems | 135 318 | 1:500 |
| mouse anti-vGlut2 | Novus Biologicals | NBP2-94571 | 1:500 |
| mouse anti-Gad67 (clone 1G10.2) | EMD Millipore | MAB5406MI | 1:500 |
| rabbit anti-synapsin1/2 | Synaptic Systems | 106 002 | 1:1000 |
| BODIPY™ 493/503 | Invitrogen | D3922 | 1ug/mL |
| mouse anti-CD43 | R&D Systems | MAB2038 | 1:100 |
| rabbit anti-PU.1 | Cell Signaling | 2266 | 1:100 |
| goat anti-Iba1 | Fisher Scientific | PIPA518488 | 1:100 |
| rabbit anti- P2RY12 | Sigma | HPA014518 | 1:50 |
| goat anti-hTrem2 | R&D Systems | AF1828 | 1:50 |
| rabbit anti-AIP1/Alix | EMD Millipore | ABC40 | 1:1000 |
| rabbit anti-Flotillin-1 | Sigma | F1180 | 1:1000 |
| Alexa Fluor 488 donkey anti-mouse IgG | Invitrogen | A21202 | 1:1000 |
| Alexa Fluor 488 goat anti-rabbit IgG | Invitrogen | A11008 | 1:1000 |
| Alexa Fluor 568 goat anti-rabbit IgG | Invitrogen | A11036 | 1:1000 |
| Alexa Fluor 568 goat anti-mouse IgG | Invitrogen | A11031 | 1:1000 |
| Alexa Fluor 647 goat anti-guinea pig IgG | Invitrogen | A21450 | 1:1000 |
| Alexa Fluor 647 donkey anti-mouse IgG | Invitrogen | A31571 | 1:1000 |
| Alexa Fluor 647 donkey anti-rabbit IgG | Invitrogen | A31573 | 1:1000 |
| Goat Anti-Rabbit IgG (H + L)-HRP Conjugate | Bio-Rad | 1706515 | 1:1000 |

**Table S3.** Reagents.

| **Reagent** | **Manufacturer** | **Catalog #** |
| --- | --- | --- |
| **iPSCs** |  |  |
| CELLSTAR 6-well cell culture plates, TC treated | Greiner Bio-One | 657160 |
| CELLSTAR 12-well cell culture plates, TC treated | Greiner Bio-One | 665102 |
| Human stem cell nucleofector kit | Lonza | VPH-5012 |
| PCR clean up kit | Qiagen | 28104 |
| mTeSR Plus | Stem Cell Technologies | 100-0276 |
| Matrigel (hESC-Qualified Matrix) | Corning | 354277 |
| Y-27632 | Stem Cell Technologies | 72304 |
| EDTA | Invitrogen | 15575-020 |
| Accutase | Stem Cell Technologies | 7920 |
| Gene JET Genomic DNA Purification kit | Thermo Fisher Scientific | K0721 |
| ATCC Universal Mycoplasma Detection Kit | ATCC | 30-1012K |
| Penicillin-Streptomycin (100X) | Life Technologies | 15140-122 |
| **Lentivirus production** |  |  |
| T175 Flasks | Greiner Bio-One | 660175 |
| Lenti-X Concentrator | Takara | 631232 |
| DMEM/F12 | Thermo Fisher Scientific | 11330-032 |
| Poly-L-Lysine | Sigma | P8920 |
| 0.45µm PVDF syringe filters | EMD Millipore | SLHVR33RS |
| **Classic NGN2 Neurons** |  |  |
| Poly-D-Lysine | Thermo Fisher Scientific | A3890401 |
| Neurobasal A | Thermo Fisher Scientific | 10888022 |
| B27 | Thermo Fisher Scientific | 17504044 |
| Non-essential amino acids | Thermo Fisher Scientific | 35050061 |
| Glutamax | Thermo Fisher Scientific | 11140050 |
| Puromycin | Gibco | A11138-03 |
| Doxycycline | Stem Cell Technologies | 72742 |
| BrdU (5-Bromo-2'-Deoxyuridine) | Thermo Fisher Scientific | B23151 |
| BDNF | PeproTech | 450-02 |
| GDNF | PeproTech | 450-10 |
| NT-3 | PeproTech | 450-03 |
| Laminin | Sigma | L2020 |
| **Enhanced NGN2 Neurons** |  |  |
| Neurobasal | Thermo Fisher Scientific | 21103049 |
| DMEM/F12 | Thermo Fisher Scientific | 11330-032 |
| Fetal bovine serum | Atlas Biologicals | F-0500-DR |
| Sodium Pyruvate | Thermo Fisher Scientific | 11360070 |
| N-Acetyl-L- cysteine | Sigma-Aldrich | A8199 |
| Heparin-Binding EGF-like Growth Factor | Sigma-Aldrich | E4643 |
| FGF | PeproTech | 100-18B |
| CNTF | PeproTech | 450-13 |
| BMP4 | PeproTech | 120-05ET |
| cAMP | Sigma | D0627 |
| KLH45 | Cayman Chemical Company | 19889 |
| **iMGLs** |  |  |
| Hematopoietic differentiation kit | Stem Cell Technologies | 5310 |
| Bambanker | Bulldog Bio | BB02 |
| DMEM/F12 (no phenol-red) | Thermo Fisher Scientific | 11039021 |
| Insulin-Transferrin-Selenium (ITS-G) 100X | Thermo Fisher Scientific | 41400045 |
| N2 | Thermo Fisher Scientific | 17502048 |
| 1-Thioglycerol (Monothioglycerol) | Sigma | M6145 |
| Human Insulin | Sigma | I9278 |
| IL-34 | PeproTech | 200-34 |
| TGF-β1 | PeproTech | 100-21 |
| M-CSF | PeproTech | 300-25 |
| CD200 | R&D systems | 2724-CD-050 |
| CX3CL1 | PeproTech | 300-31 |
| Lipopolysaccharide (LPS) | Sigma-Aldrich | L4516 |
| Latex beads, amine-modified polystyrene, fluorescent red | Sigma-Aldrich | L2778 |
| **RT-qPCR** |  |  |
| Directzol RNA MicroPreps | Zymo Research | NC1000412 |
| SuperScript IV First Strand Synthesis System | Thermo Fisher Scientific | 18091050 |
| iTaq Univer SYBR Green Supermix | Bio-Rad | 1725122 |
| **Ca-Imaging** |  |  |
| Fluo-4AM | Thermo Fisher Scientific | F14201 |
| DMSO | Thermo Fisher Scientific | BP231100 |
| Pluronic acid 20% solution in DMSO | Thermo Fisher Scientific | P3000MP |
| NaCl | Fisher Bioreagents | BP358-212 |
| KCl | Sigma-Aldrich | P4504 |
| CaCl<sub>2</sub> | Sigma-Aldrich | C-3881 |
| MgCl<sub>2</sub> | VWR | E525 |
| HEPES | Biopioneer | C0113 |
| Glucose | Sigma | G5400 |
| **Immunocytochemistry** |  |  |
| Glass-bottom 96-well plates, #1.5 cover glass | Cellvis | NC0536760 |
| 16% Paraformaldehyde solution | Electron Microscopy Sciences | 50980487 |
| PBS | Fisher | 14190 |
| Bovine serum albumin | Sigma | A7906 |
| TritonX-100 | Sigma | T9284 |
| **EV Isolation** |  |  |
| SW32 Ti Swinging-Bucket Rotor | Beckman Coulter | 369650 |
| Polypropylene centrifuge tubes | Beckman Coulter | 326823 |
| Ultra-clear centrifuge tubes | Beckman Coulter | 344058 |
| DPBS, calcium, magnesium | Life Technologies | 14040117 |
| DC Protein Assay | Bio-Rad | 5000112 |
| **Western Blot** |  |  |
| Protease Inhibitor Tablets | Thermo Fisher Scientific | A32955 |
| 6X Laemmli SDS sample buffer | Thermo Fisher Scientific | J61337AC |
| Trans-Blot Turbo Mini NC Transfer Packs | Bio-Rad | 1704158 |
| 4-20% MP TGX Gel 10W 50 µl | Bio-Rad | 4561094 |
| 10X Tris/Glycine Buffer | Bio-Rad | 1610734 |
| Super Signal West Femto Maximum Sensitivity Substrate | Thermo Fisher Scientific | 34095 |
| Ponceau S solution | Sigma-Aldrich | 9189 |
| Tween20 | Fisher | BP337-100 |
| Tris-HCl pH 7.5 | Invitrogen | 15567-027 |

**Table S4.** Median and Quartile Data for Violin Plots in Main Figures.

| Figure | Panel | Median, 25%, and 75% quartiles |
| --- | --- | --- |
| 1 | I | PBS, 1.00, 0.83, 1.24; PBS-Exos, 2.07, 1.00, 2.90; LPS-Exos, 1.50, 1.25, 2.49 |
| 1 | J | PBS, 0.92, 0.75, 1.17; PBS-Exos, 0.94, 0.77, 1.17; LPS-Exos, 0.98, 0.78, 1.24 |
| 1 | K | PBS, 6, 2.75, 8.75; PBS-Exos, 7, 6, 8.25; LPS-Exos, 10, 6.5, 13 |
| 1 | L | PBS, 1, 0.25, 2.75; PBS-Exos, 15.5, 3.75, 7; LPS-Exos, 5, 4, 8.5 |
| 1 | M | PBS, 0.71, 0.64, 0.77; PBS-Exos, 0.88, 0.78, 0.97; LPS-Exos, 0.84, 0.76, 0.89 |
| 1 | N | PBS, 0.60, 0.56, 0.66; PBS-Exos, 0.67, 0.59, 0.77; LPS-Exos, 0.65, 0.57, 0.72 |
| 2 | C | PBS UCM, 0.84, 0.56, 1.21; E3 PBS CM, 0.78, 0.54, 1.12; E4 PBS CM, 0.86, 0.61, 1.21; LPS UCM, 0.89, 0.65, 1.23; E3 LPS CM, 0.97, 0.70, 1.32; E4 LPS CM, 1.14, 0.79, 1.51 |
| 2 | D | PBS UCM, 0.70, 0.66, 1.32; E3 PBS CM, 0.68, 0.68, 1.36; E4 PBS CM, 1.16, 0.68, 1.39; LPS UCM, 0.61, 0.61, 1.21; E3 LPS CM, 1.52, 0.98, 2.37; E4 LPS CM, 2.29, 1.32, 3.94 |
| 2 | E | PBS UCM, 0, 0, 1; E3 PBS CM, 0, 0, 1; E4 PBS CM, 0.5, 0, 2; LPS UCM, 0, 0, 1; E3 LPS CM, 3, 2, 5; E4 LPS CM, 5, 0.75, 8 |
| 2 | F | PBS UCM, 0, 0, 0; E3 PBS CM, 0, 0, 0; E4 PBS CM, 0, 0, 0; LPS UCM, 0, 0, 0; E3 LPS CM, 1, 0, 3.75; E4 LPS CM, 3.5, 0, 6 |
| 2 | H | PBS UCM, 0.88, 0.70, 1.14; E3 PBS CM, 0.97, 0.72, 1.33; E4 PBS CM, 0.87, 0.68, 1.19; LPS UCM, 0.86, 0.71, 1.16; E3 LPS CM, 1.16, 0.89, 1.59; E4 LPS CM, 1.23, 0.90, 1.68 |
| 2 | I | PBS UCM, 0.88, 0.45, 1.24; E3 PBS CM, 0.90, 0.50, 1.77; E4 PBS CM, 0.88, 0.50, 1.35; LPS UCM, 0.60, 0.59, 1.2; E3 LPS CM, 1.51, 1.10, 2.93; E4 LPS CM, 2.72, 1.82, 5.85 |
| 2 | J | PBS UCM, 0, 0, 1; E3 PBS CM, 0, 0, 0.75; E4 PBS CM, 0, 0, 1; LPS UCM, 0, 0, 0.75; E3 LPS CM, 4, 0.75, 6.25; E4 LPS CM, 8, 2.75, 11.25 |
| 2 | K | PBS UCM, 0, 0, 0; E3 PBS CM, 0, 0, 0; E4 PBS CM, 0, 0, 0; LPS UCM, 0, 0, 0; E3 LPS CM, 1, 0, 4; E4 LPS CM, 4, 0, 7 |
| 2 | L | E3CM/E3iNs, 0.97, 0.70, 1.32; E3CM/E4iNs, 1.16, 0.89, 1.59; E4CM/E3iNs, 1.14, 0.79, 1.51; E4CM/ E4iNs, 1.23, 0.90, 1.68 |
| 2 | M | E3CM/E3iNs, 1.52, 0.98, 2.37; E3CM/E4iNs, 1.51, 1.10, 2.93; E4CM/E3iNs, 2.29, 1.32, 3.94; E4CM/ E4iNs, 2.72, 1.82, 5.8 |
| 2 | N | E3CM/E3iNs, 3, 2, 5; E3CM/E4iNs, 4, 0.75, 6.25; E4CM/E3iNs, 5, 0.75, 8; E4CM/ E4iNs, 8, 2.75, 11.25 |
| 2 | O | E3CM/E3iNs, 1, 0, 3.75; E3CM/E4iNs, 1, 0, 4; E4CM/E3iNs, 3.5, 0, 6; E4CM/ E4iNs, 4, 0, 7 |
| 4 | H | Veh, 0.95, 0.73, 1.28; KLH45 Pre, 0.50, 0.25, 0.76; KLH45 Pre+Post, 0.30, 0.10, 0.78 |
| 4 | I | Veh, 0.90, 0.69, 1.20; KLH45 Pre, 0.57, 0.46, 0.72; KLH45 Pre+Post, 0.54, 0.42, 0.72 |
| 4 | J | Veh, 7, 3, 9; KLH45 Pre, 0, 0, 2; KLH45 Pre+Post, 0, 0, 0 |
| 4 | K | Veh, 0.82, 0.49, 1.25; KLH45 Pre, 0.45, 0.20, 0.75; KLH45 Pre+Post, 0.40, 0.24, 0.82 |
| 4 | L | Veh, 0.84, 0.66, 1.16; KLH45 Pre, 0.70, 0.55, 0.94; KLH45 Pre+Post, 0.66, 0.54, 0.84 |
| 4 | M | Veh, 3, 1, 5; KLH45 Pre, 0, 0, 0; KLH45 Pre+Post, 1, 0, 2 |

## References

1. S. Marinelli, B. Basilico, M. C. Marrone, D. Ragozzino, Microglia-neuron crosstalk: Signaling mechanism and control of synaptic transmission. Semin Cell Dev Biol 94, 138–151 (2019).

2. R. Akiyoshi et al., Microglia Enhance Synapse Activity to Promote Local Network Synchronization. eneuro 5, ENEURO.0088-0018.2018 (2018).

3. A. A.-O. X. Clark et al., Selective activation of microglia facilitates synaptic strength. (2015).

4. J.-C. Lambert et al., Meta-analysis of 74,046 individuals identifies 11 new susceptibility loci for Alzheimer’s disease. Nature Genetics 45, 1452–1458 (2013).

5. Celeste M. Karch, C. Cruchaga, Alison M. Goate, Alzheimer’s Disease Genetics: From the Bench to the Clinic. Neuron 83, 11–26 (2014).

6. D. P. Wightman et al., A genome-wide association study with 1,126,563 individuals identifies new risk loci for Alzheimer’s disease. Nature Genetics 53, 1276–1282 (2021).

7. B. W. Kunkle et al., Genetic meta-analysis of diagnosed Alzheimer’s disease identifies new risk loci and implicates Aβ, tau, immunity and lipid processing. Nature Genetics 51, 414–430 (2019).

8. U. B. Eyo et al., Regulation of Physical Microglia-Neuron Interactions by Fractalkine Signaling after Status Epilepticus. eNeuro 3 (2016).

9. K. M. Rodgers et al., The cortical innate immune response increases local neuronal excitability leading to seizures. Brain 132, 2478–2486 (2009).

10. B. Chausse et al., Metabolic flexibility ensures proper neuronal network function in moderate neuroinflammation. Sci Rep 14, 14405 (2024).

11. G. H. Park et al., Activated microglia cause metabolic disruptions in developmental cortical interneurons that persist in interneurons from individuals with schizophrenia. Nat Neurosci 23, 1352–1364 (2020).

12. M. Mathieu, L. Martin-Jaular, G. Lavieu, C. Thery, Specificities of secretion and uptake of exosomes and other extracellular vesicles for cell-to-cell communication. Nat Cell Biol 21, 9–17 (2019).

13. J. Li et al., Exosomes in Central Nervous System Diseases: A Comprehensive Review of Emerging Research and Clinical Frontiers. 10.3390/biom14121519.

14. Y.-F. Chen, F. Luh, Y.-S. Ho, Y. Yen, Exosomes: a review of biologic function, diagnostic and targeted therapy applications, and clinical trials. Journal of Biomedical Science 31, 67 (2024).

15. Y. Yang et al., Inflammation leads to distinct populations of extracellular vesicles from microglia. Journal of Neuroinflammation 15, 168 (2018).

16. J. V. Santiago et al., Identification of State-Specific Proteomic and Transcriptomic Signatures of Microglia-Derived Extracellular Vesicles. Mol Cell Proteomics 22, 100678 (2023).

17. V. Budnik, C. Ruiz-Canada, F. Wendler, Extracellular vesicles round off communication in the nervous system. Nat Rev Neurosci 17, 160–172 (2016).

18. G. C. De Paula et al., Extracellular vesicles released from microglia after palmitate exposure impact brain function. J Neuroinflammation 21, 173 (2024).

19. C. Fan et al., Microglia secrete miR-146a-5p-containing exosomes to regulate neurogenesis in depression. Mol Ther 30, 1300–1314 (2022).

20. X. Liu et al., New insights on targeting extracellular vesicle release by GW4869 to modulate lipopolysaccharide-induced neuroinflammation in mice model. Nanomedicine (Lond) 19, 2619–2632 (2024).

21. E. H. Corder et al., Gene dose of apolipoprotein E type 4 allele and the risk of Alzheimer’s disease in late onset families. Science 261, 921–923 (1993).

22. M. S. Haney et al., APOE4/4 is linked to damaging lipid droplets in Alzheimer’s disease microglia. Nature 628, 154–161 (2024).

23. M. P. Vitek, C. M. Brown, C. A. Colton, APOE genotype-specific differences in the innate immune response. Neurobiology of Aging 30, 1350–1360 (2009).

24. J. R. Lynch et al., APOE genotype and an ApoE-mimetic peptide modify the systemic and central nervous system inflammatory response. (2003).

25. Y. T. Lin et al., APOE4 Causes Widespread Molecular and Cellular Alterations Associated with Alzheimer’s Disease Phenotypes in Human iPSC-Derived Brain Cell Types. Neuron 98, 1141–1154 e1147 (2018).

26. C.-C. Liu et al., Cell-autonomous effects of APOE4 in restricting microglial response in brain homeostasis and Alzheimer’s disease. Nature Immunology 24, 1854–1866 (2023).

27. M. B. Victor et al., Lipid accumulation induced by APOE4 impairs microglial surveillance of neuronal-network activity. Cell Stem Cell 29, 1197–1212 e1198 (2022).

28. A. Rao et al., Microglia depletion reduces human neuronal APOE4-related pathologies in a chimeric Alzheimer’s disease model. Cell Stem Cell 32, 86–104 e107 (2025).

29. D. R. Tabuena et al., Neuronal APOE4-induced Early Hippocampal Network Hyperexcitability in Alzheimer’s Disease Pathogenesis. bioRxiv 10.1101/2023.08.28.555153 (2024).

30. S. Raman, N. Brookhouser, D. A. Brafman, Using human induced pluripotent stem cells (hiPSCs) to investigate the mechanisms by which Apolipoprotein E (APOE) contributes to Alzheimer’s disease (AD) risk. Neurobiol Dis 138, 104788 (2020).

31. V. Lo Sardo et al., Unveiling the Role of the Most Impactful Cardiovascular Risk Locus through Haplotype Editing. Cell 175, 1796–1810 e1720 (2018).

32. Y. Zhang et al., Rapid single-step induction of functional neurons from human pluripotent stem cells. Neuron 78, 785–798 (2013).

33. I. Canals et al., Rapid and efficient induction of functional astrocytes from human pluripotent stem cells. Nat Methods 15, 693–696 (2018).

34. A. McQuade et al., Development and validation of a simplified method to generate human microglia from pluripotent stem cells. Mol Neurodegener 13, 67 (2018).

35. R. A. Stephenson et al., Triglyceride metabolism controls inflammation and APOE4-associated disease states in microglia. bioRxiv 10.1101/2024.04.11.589145 (2024).

36. G. Sienski et al., APOE4 disrupts intracellular lipid homeostasis in human iPSC-derived glia. Sci Transl Med 13 (2021).

37. P. Sharma et al., Exosomes regulate neurogenesis and circuit assembly. Proc Natl Acad Sci U S A 116, 16086–16094 (2019).

38. J. A. Olzmann, P. Carvalho, Dynamics and functions of lipid droplets. Nat Rev Mol Cell Biol 20, 137–155 (2019).

39. M. Kumar et al., DDHD2 is necessary for activity-driven fatty acid fueling of nerve terminal function. BioRXIV preprint 10.1101/2023.12.18.572201 (2023).

40. J. M. Inloes et al., The hereditary spastic paraplegia-related enzyme DDHD2 is a principal brain triglyceride lipase. Proceedings of the National Academy of Sciences 111, 14924–14929 (2014).

41. O. Pascual, S. Ben Achour, P. Rostaing, A. Triller, A. Bessis, Microglia activation triggers astrocyte-mediated modulation of excitatory neurotransmission. Proc Natl Acad Sci U S A 109, E197–205 (2012).

42. U. B. Eyo, M. Murugan, L. J. Wu, Microglia-Neuron Communication in Epilepsy. Glia 65, 5–18 (2017).

43. F. Antonucci et al., Microvesicles released from microglia stimulate synaptic activity via enhanced sphingolipid metabolism. EMBO J 31, 1231–1240 (2012).

44. J. Marschallinger et al., Lipid-droplet-accumulating microglia represent a dysfunctional and proinflammatory state in the aging brain. Nat Neurosci 23, 194–208 (2020).

45. S. Zhao et al., Chemogenetic activation of microglial Gi signaling decreases microglial surveillance and impairs neuronal synchronization. Science Advances 11, eado7829 (2025).

46. N. Filippini et al., Distinct patterns of brain activity in young carriers of the APOE-ε4 allele. Proceedings of the National Academy of Sciences 106, 7209–7214 (2009).

47. T. Nuriel et al., Neuronal hyperactivity due to loss of inhibitory tone in APOE4 mice lacking Alzheimer’s disease-like pathology. Nat Commun 8, 1464 (2017).

48. K. Mann, S. Deny, S. Ganguli, T. R. Clandinin, Coupling of activity, metabolism and behaviour across the Drosophila brain. Nature 593, 244–248 (2021).

49. O. Stoler et al., Frequency- and spike-timing-dependent mitochondrial Ca2+ signaling regulates the metabolic rate and synaptic efficacy in cortical neurons. eLife 11, e74606 (2022).

50. Janneke H. M. Schuurs-Hoeijmakers et al., Mutations in DDHD2, Encoding an Intracellular Phospholipase A1, Cause a Recessive Form of Complex Hereditary Spastic Paraplegia. The American Journal of Human Genetics 91, 1073–1081 (2012).

51. J. M. Inloes et al., Functional Contribution of the Spastic Paraplegia-Related Triglyceride Hydrolase DDHD2 to the Formation and Content of Lipid Droplets. Biochemistry 57, 827–838 (2018).

52. I. O. Akefe et al., The DDHD2-STXBP1 interaction mediates long-term memory via generation of saturated free fatty acids. The EMBO Journal 43, 533–567-567 (2024).

53. C. Yang et al., Rewiring Neuronal Glycerolipid Metabolism Determines the Extent of Axon Regeneration. Neuron 105, 276–292 e275 (2020).

54. I. Ralhan, C. L. Chang, J. Lippincott-Schwartz, M. S. Ioannou, Lipid droplets in the nervous system. J Cell Biol 220 (2021).

## SI References

1. V. Lo Sardo et al., Unveiling the Role of the Most Impactful Cardiovascular Risk Locus through Haplotype Editing. Cell 175, 1796–1810 e1720 (2018).

2. A. McQuade, M. Blurton-Jones, “Human Induced Pluripotent Stem Cell-Derived Microglia (hiPSC-Microglia)” in Induced Pluripotent Stem (iPS) Cells: Methods and Protocols, A. Nagy, K. Turksen, Eds. (Springer US, New York, NY, 2022), 10.1007/7651_2021_429, pp. 473–482.

3. X. Qiu et al., Distinct functions of dimeric and monomeric scaffold protein Alix in regulating F-actin assembly and loading of exosomal cargo. Journal of Biological Chemistry 298, 102425 (2022).

